# Immune and Mutational Profile of Gene-Edited Low-Immunogenic Human Primary Cholangiocyte Organoids

**DOI:** 10.1101/2025.01.20.628680

**Authors:** Sandra Petrus-Reurer, Adrian Baez-Ortega, Winnie Lei, Girishkumar Kumaran, James Williamson, Julia Jones, Daniel Trajkovski, Andrew R.J. Lawson, Krishnaa T. Mahbubani, Cara Brodie, Paul Lehner, Matthew J. Bottomley, Inigo Martincorena, Kourosh Saeb-Parsy

**Affiliations:** Department of Surgery, University of Cambridge and NIHR Cambridge Biomedical Research Centre, Cambridge, United Kingdom; Wellcome Sanger Institute, Wellcome Genome Campus, Hinxton, United Kingdom; CAMS Oxford Institute, Nuffield Department of Medicine, Oxford, United Kingdom; Cambridge Institute for Therapeutic Immunology and Infectious Disease, Jeffrey Cheah Biomedical Centre, Cambridge Biomedical Campus, University of Cambridge, Cambridge, UK; Cancer Research UK Cambridge Institute, Cambridge, United Kingdom Kingdom

## Abstract

Primary human cells cultured in organoid format have great promise as potential regenerative cellular therapies. However, their immunogenicity and mutational profile remain unresolved, impeding effective long-term translation to the clinic. In this study we report, for the first time, the generation of human leukocyte antigen (HLA)-I and HLA-II knock-out expandable human primary cholangiocyte organoids (PCOs) using CRISPR-Cas9 as a potential ‘universal’ low-immunogenic therapy for bile duct disorders. HLA-edited PCOs (ePCOs) displayed the same phenotypical and functional characteristics as parental un-edited PCOs. Despite minimal off-target edits, single-molecule DNA-sequencing demonstrated that ePCOs and PCOs acquire substantial mutations in culture at similar rates but without evident selection for cancer-driver mutations. ePCOs induced reduced T cell-mediated immunity and a donor-dependent NK cell cytotoxicity *in vitro* and evaded cytotoxic responses with increased graft survival in humanized mice *in vivo*. Our findings have important implications for assessment of safety and immunogenicity of organoid cellular therapies.

## INTRODUCTION

Cellular therapies hold great potential for restoring damaged or diseased cells and tissues as a consequence of aging, disease, injury or accidents^1,2^. Typically, these therapies involve deriving cell types of interest from either an undifferentiated source of cells with multi- or pluripotency (including human embryonic [hESC] or induced [hiPSC] pluripotent stem cell origin)^1,3^, or expansion of fully mature primary cells typically in a 3D cell culture format known as organoids^4^. Organoids, derived from primary rather than pluripotent cells, are a novel and promising technology for cellular therapies due to the presence of multiple tissue-specific cell types that recapitulate the normal phenotype and function better than 2D cell models^5–9^. However, the immunogenicity of these cellular products still remains an unresolved challenge that may impede effective long-term translation to the clinic^10^.

Immune response to most allogeneic solid organs is driven predominantly by human leukocyte antigen (HLA) molecules^11–13^. Hence, allogeneic cellular therapies are similarly expected to induce an immune response via direct and indirect allorecognition mechanisms driven in large part by HLA^10^. Different strategies are pursued to reduce the rejection of allogeneic ‘off-the-shelf’ cellular therapies, including donor-recipient matching, use of classical immunosuppressive drugs, use of cell shielding materials, or use of T regulatory cells^10^. Another strategy to evade the potential immune rejection of donor cells is the generation of allogeneic ‘universal’ therapies by genetic engineering to abolish expression of HLA Class I and II. Although multiple studies have shown that pluripotent stem cells (hESC/hiPSC lines) can be genetically manipulated for this purpose^14^, the generation and subsequent comprehensive immunogenicity evaluation of low-immunogenic HLA-edited human primary proliferative organoid systems has not yet been achieved.

The assessment of off-target mutations after CRISPR-Cas9 editing is a critical step for the evaluation of safety of the cellular product as established by several studies^15–17^. However, culturing and passaging cells can also introduce mutations that could pose safety concerns as previously reported^18–20^. Mutational analysis of polyclonal cell populations using standard whole-genome or targeted sequencing can only detect mutations that affect a substantial proportion of cells, which complicates the screening of oncogenic mutations in cell therapy lines^20^. However, new technologies such as nanorate sequencing (NanoSeq)^21^, a duplex sequencing method that enables the accurate detection and quantification of mutations present in single DNA molecules, can circumvent these limitations to support sensitive mutational safety assessments of polyclonal lines^22^. The incorporation of these novel technologies in the safety evaluation of both edited and unedited organoid cell therapies could considerably de-risk their clinical translation.

We have previously shown that primary cholangiocyte organoids (PCOs) can be derived from human cholangiocytes without genetic editing and have potential for use as cellular and bioengineered therapies for bile duct disorders^23–25^, which currently have no curative treatment other than complex surgery or transplantation. In this study we efficiently engineered HLA-I and II double knock-out (DKO) human PCOs (ePCOs) using a CRISPR-Cas9 system. ePCOs displayed unaltered morphology, phenotype and functionality compared to wild-type cells. We also show that editing induced minimal off-target mutations, none of which affects cancer genes. Importantly, NanoSeq revealed that passaging in culture is a much more substantial source of mutations than CRISPR-Cas9 edits, although we did not detect evidence of cancer-driver mutations in our lines. Upon *in vitro* co-culture with allogeneic human immune cells, ePCOs induced significantly reduced immune activation, although NK cells showed donor-dependent cytotoxic activity towards both parental and engineered cells. In a humanized mouse model, ePCOs showed better preserved graft survival and significantly reduced local immune infiltration compared to parental unedited controls, due to evasion of T cell mediated cytotoxic responses and downregulated cell graft stress and extrinsic apoptotic pathways. This work provides the first report on the generation of HLA-edited expandable human primary organoids and a high-resolution analysis of their mutational profile and immunogenicity with state-of-the-art technologies and in relevant experimental *in vitro* and *in vivo* systems.

## RESULTS

### HLA DKO ePCO lines can be efficiently generated using CRISPR-Cas9 system targeting *B2M* and *RFX5* loci

Three lines of human primary-derived cholangiocyte organoids (**Supplementary Table 1**) were edited from parental donor lines (parental PCOs) by two sequential rounds of electroporation with CRISPR-Cas9 RNP molecules with guides targeting coding regions of *B2M* and *RFX5* genes (**Supplementary Table 2**). Cells were further plated, expanded and stimulated with IFN-ɣ and subsequently sorted using flow cytometry for HLA-I and HLA-II double negative populations (DKO ePCO) with efficiencies after sort of 58-89% (**Fig. 1a, Extended Data Fig. 1a**). After sorting, single cells were plated and expanded for characterisation. ePCOs showed the same sphere-like morphology and expression of CD326 marker as parental cells. Importantly, even when cultured with IFN-ɣ to simulate pro-inflammatory conditions, ePCOs lacked HLA-I and HLA-II expression as expected (**Fig. 1b, Extended Data Fig. 1b**). Functionally, ePCOs displayed similar levels of specific enzymatic activity to both parental and primary tissue, as measured by gamma glutamyl transferase (GGT) assay and alkaline phosphatase (ALP) staining, with activity similar to unedited cholangiocyte organoids in previous studies^23,24^ (**Fig. 1c**). One line (ePCO line 2) was advanced to more detailed proteomic analysis and *in vivo* studies. Proteomic analysis confirmed that HLA-related and HFE (which regulates production of hepcidin, binding to transferrin receptor) proteins were the only differentially expressed species on surface of ePCOs (**Fig. 1d**, **Extended Data Fig. 1c, Supplementary Table 3**). After injection under kidney capsule of immunocompromised MHC-I/II dKO NOD scid gamma (NSG) mice, ePCO successfully engrafted and expressed specific human and cholangiocyte markers (hKu80, hKRT7) at 2 weeks and 5 weeks after injection with similar engraftment pattern as parental unedited organoids (**Fig. 1e**).

**FIG. 1.**
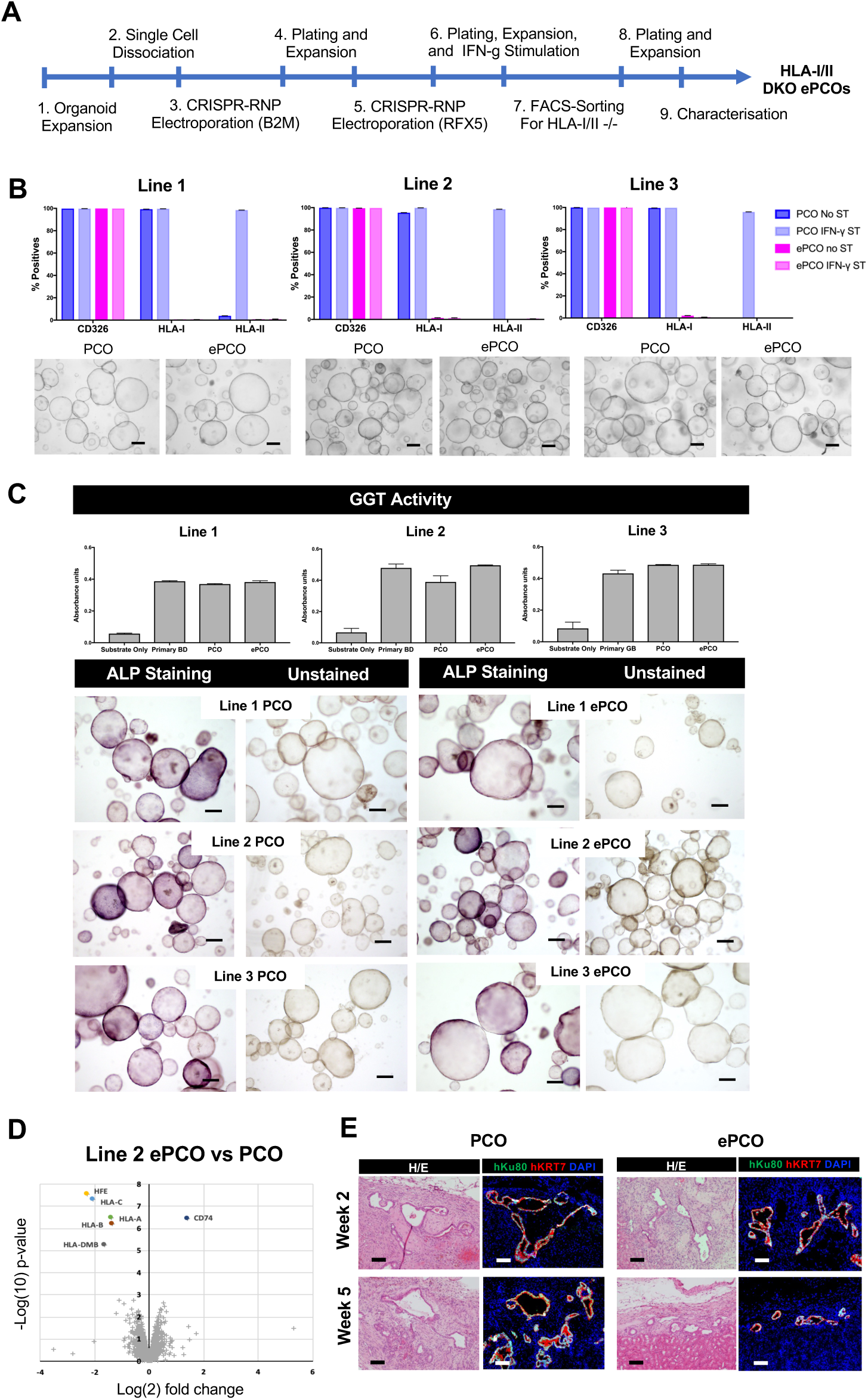
Generation and characterisation of DKO ePCOs. **(a)** Diagram of the process used to generate HLA-I/II DKO ePCOs. (**b)** Bar graphs showing percentage of cell surface expression of CD326, HLA-I and HLA-II by flow cytometry for PCO and ePCO organoids cultured with and without IFN-ɣ stimulation (2 days, 100ng/mL) for the three PCO/ePCO lines. Fluorescence minus one (FMO) controls are used for gating (top). Bright field pictures of PCO and ePCO cells for the three lines are shown (bottom). Error bars represent mean±SEM from three independent experiments. Scale-bars: 200 µm. **(c)** Bar graphs showing gamma glutamyl transferase activity (GGT) for PCO, ePCO and respective primary tissue. Negative control is substrate only (top). Bright field pictures showing alkaline phosphatase (ALP) staining in PCO and respective ePCO organoids. Negative control is unstained (bottom). Scale-bars: 200 µm. **(d)** Plot showing differentially expressed surface proteins in Line 2 ePCO compared to its parental PCO. Cut-off q-value of <=0.01 and fold change cut off of 2-fold. **(e)** Haematoxylin/Eosin and immunofluorescence staining showing PCO and ePCO engraftment, and specific human Ku80 and KRT7 staining at 2 and 5 weeks after injection of PCO and ePCO under kidney capsule. Scale-bars: 100 µm.

### Genetic editing induces minimal off-target mutations not overlapping with cancer genes

In order to assess CRISPR on-target specificity we analysed rates, patterns and impacts of mutation accumulation in ePCOs. To quantify mutations, we sequenced the genomes of primary tissue, DKO and parental organoids at different passages (P0, P1, P5) from the three ePCO lines (**Supplementary Table 4**), using standard whole-genome sequencing (WGS) and nanorate sequencing (NanoSeq)^21^ (**Fig. 2a**). WGS analysis confirmed the presence of complete CRISPR-mediated knock-out of genes *B2M* and *RFX5* in ePCO samples from all three lines (**Extended Data Fig. 2a**). Despite the potential for protein-coding regions to be affected by a small fraction of predicted CRISPR off-target sites, identification of mutations through WGS revealed virtually no high-frequency mutations in these regions in any of the three ePCO lines compared to respective parental unedited samples (**Table 1**). Additionally, 5-17 non-synonymous mutations were detected in ePCO lines outside predicted off-target sites, none of which were listed in the COSMIC Cancer Gene Sensus^26^ (**Supplementary Table 5**). Except for deletions at the two target loci, mutations did not display spatial clustering or other signs associated with CRISPR editing (see **Methods**). These results demonstrate that generation of ePCOs via CRISPR did not produce evident alterations in non-targeted protein-coding genes, including cancer-associated genes. Additionally, no chromosomal aberrations were detected in ePCO Lines 1 and 2. However, a chromosome 7 trisomy was detected in ePCO line 3 (**Fig. 2b, Extended Data Fig. 2b**). This is likely a spontaneous consequence of cell culture (as previously reported for other cells in culture^27,28^), or a preexisting somatic event expanded *in vitro* (as somatic chromosome 7 trisomies have been reported with age in some tissues^29^), rather than the result of CRISPR editing. Nonetheless, this result highlights the need for genomic screening of future engineered lines, as similar aberrations would preclude them from being advanced towards the clinic.

**FIG. 2.**
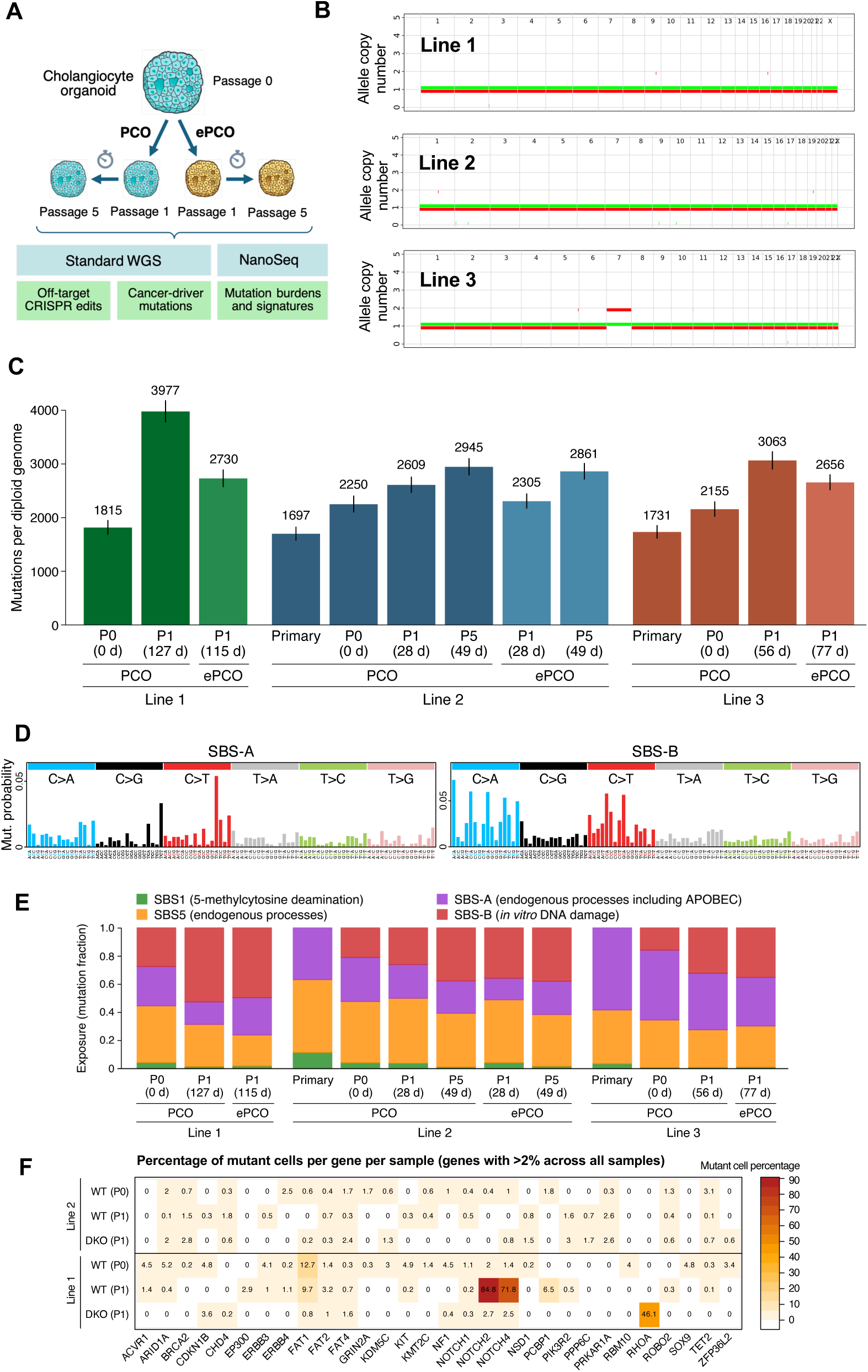
Mutation burden in ePCOs. **(a)** Schematic diagram of the experimental and sequencing design. **(b)** Diagram of allele-specific copy number along the genome for each engineered line, with alleles shown in different colours. **(c)** Mutation burdens per cell (diploid genome) as estimated through NanoSeq in each sample. Number of mutations per genome is indicated above each bar. Error bars represent 95% confidence intervals. Samples from each line are shown in a different colour, with DKO samples coloured in a lighter shade. Passage number (P) and days in culture (d) are indicated for each sample. **(d)** Mutational spectra of the two signatures inferred *de novo* (SBS-A; SBS-B). **(e)** Estimated mutational exposures (fractions of mutations per signature) for four identified mutational signatures in each sample. **(f)** Estimated percentage of cells carrying at least one coding gene mutation in three organoid samples (WT P0, WT P1, DKO P1) from PCO lines 2 and 3. Only genes with estimated mutant cell percentages greater than 2% are shown.

**TABLE 1.** Mutational analysis of predicted CRISPR off-target sites. Table summarising pathogenicity analysis of predicted CRISPR off-target sites, including numbers (percentages) of predicted off-target sites overlapping coding sequences (CDS), and numbers (percentages) of mutations in predicted off-target sites as identified through WGS in each line.

|  |  |  |
| --- | --- | --- |
| **Off-target sites overlapping protein-coding gene CDS** |  | 2218 (3.67%) |
| **Off-target sites overlapping Cancer Gene Census CDS** |  | 92 (0.15%) |
| **Mutations overlapping any off-target site** | Line 1 | 5 (0.28%) |
|  | Line 2 | 0 (0.00%) |
|  | Line 3 | 3 (0.17%) |
| **Mutations overlapping protein-coding CDS in off-target sites** | Line 1 | 0 (0.00%) |
|  | Line 2 | 0 (0.00%) |
|  | Line 3 | 1 (0.06%) |
| **Mutations overlapping Cancer Gene Census CDS in off-target sites** | Line 1 | 0 (0.00%) |
|  | Line 2 | 0 (0.00%) |
|  | Line 3 | 0 (0.00%) |

### NanoSeq enables sensitive assessment of mutation accumulation and clonal expansion *in vitro*

Although concerns about CRISPR off-target edits have dominated discussions around the safety of some cellular therapies^16,30,31^, *in vitro* culture is itself mutagenic and can cause considerable numbers of mutations in single cells during cell culture^18,19^. This includes potentially oncogenic mutations, some of which can expand during culture and passage^20^. Monitoring these mutations is important for safety considerations, but they can be undetectable in polyclonal cultures with standard sequencing methods. Here we used NanoSeq to estimate the mutation burden per cell for each ePCO sample^21,22^. This provided evidence for progressive acquisition of mutations during culture in all lines (**Fig. 2c**). We employed a Bayesian statistical approach^32^ to deconvolute the mutations in each ePCO sample into a set of mutational signatures. Each of these signatures represents one or more mutational processes that have contributed mutations at some point between the donor’s fertilised zygote and the somatic cells in each ePCO sample^33^. Specifically, we identified four mutational signatures, including two well-known signatures (SBS1 and SBS5) that reflect spontaneous deamination of 5-methylcytosine and other ubiquitous endogenous mutational processes, characteristic of age-related *in vivo* somatic mutagenesis in most human tissues^34^; and two *de novo* signatures, labelled SBS-A and SBS-B (**Fig. 2d-e**). SBS-A was detectable in all samples, including ePCOs and pre-culture primary tissue samples, and likely represents a combination of *in vivo* endogenous mutational processes, possibly including APOBEC activity (characterised by C>T and C>G mutations at TCN sequence contexts^34^). In contrast, SBS-B was specific to ePCO samples and tended to increase in prominence over time in culture (**Fig. 2e**), suggesting that this signature represents continuous DNA damage *in vitro*. This type of damage has been shown to be dominated by oxidative damage due to reactive oxygen species, characterised by a high rate of C>A mutations^35–37^. Overall, these results suggest that *in vitro* culture is a much larger source of new mutations than CRISPR off-target edits, requiring careful monitoring for cellular therapies.

Beyond an overall increase of mutations per cell, the occurrence of cancer-related mutations and the possibility that organoid derivation or tissue culture may select oncogenic mutations could pose a safety risk for clinical use of ePCOs. In polyclonal cultures, these mutations can exist at allele frequencies below the detection sensitivity of standard DNA sequencing methods (which is typically ∼1-5%). To more sensitively screen for oncogenic mutations, we performed targeted NanoSeq on a panel of 257 cancer genes (median duplex depth, 650x). This analysis did not reveal convincing evidence of positive selection of cancer-driver mutations in our ePCO lines (**Supplementary Table 6**). Although three clonal expansions in P1 relative to P0 had one mutation in a cancer driver gene (*NOTCH2*, *NOTCH4*, *RHOA*) (**Fig. 2f**), these expansions appear more likely to reflect culture or passaging bottlenecks than selection (**Supplementary Table 7**).

Taken together, our results demonstrate that generation of ePCOs via CRISPR-Cas9 editing does not seem to be a significant source of mutations compared to unedited controls. Importantly, no off-target mutations overlapped with cancer genes. However, we show that both parental and edited cells accumulate a much larger number of additional mutations due to DNA damage during culture, consistent with previous studies^35–37^. As these mutations can escape undetected by standard sequencing methods in polyclonal cultures, sensitive technologies like NanoSeq may be needed to monitor cellular therapy lines. Reassuringly, despite *in vitro* culture inducing a considerable number of mutations per cell, we observed no convincing evidence of selection or clonal expansion of cells carrying cancer-driver mutations.

### HLA DKO ePCOs induce significantly reduced immune reactivity *in vitro*

The immunological consequence of HLA I/II deletion in ePCOs was first assessed by *in vitro* co-culture with allogenic human donor peripheral blood mononuclear cells (PBMCs). The specific organoid:PBMC ratio was first evaluated using two parental PCO lines co-cultured with different PBMC donors at 1:5, 1:10, 1:20 and 1:50 ratio. In all cases IFN-ɣ secretion was higher than in negative controls, but 1:10 ratio resulted in higher secretion levels and was therefore selected for subsequent experiments (**Extended Data Fig. 3a**). When evaluating parental PCOs and DKO ePCOs with same assay at 1:10 ratio, all three ePCO lines induced significantly reduced IFN-ɣ secretion compared to parental controls, thus demonstrating reduced activation of allogeneic T cells (**Fig. 3a**). Anti-inflammatory cytokines IL-10 and IL-6 were decreased in ePCO co-cultures in 2/3 donors, while there were no significant differences in other analyzed cytokines (**Supplementary Table 8**).

**FIG. 3.**
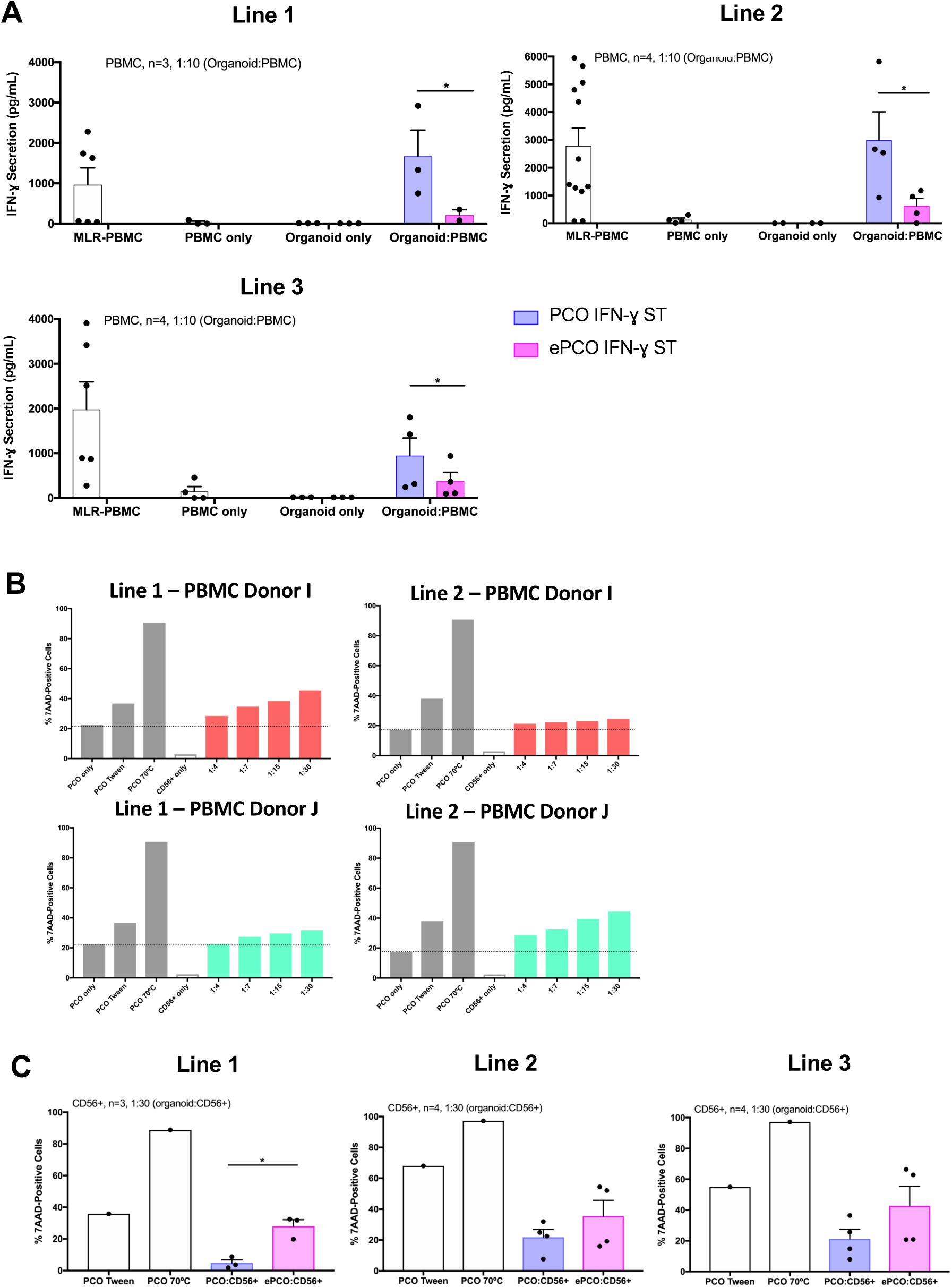
Immunogenicity of ePCOs *in vitro*. **(a)** Bar graphs showing concentration of IFN-ɣ secretion by allogeneic PBMCs when co-cultured (1:10 organoid:PBMC ratio) with IFN-ɣ pre-stimulated PCO and ePCO for 5 days. Negative controls are PCOs only and PBMCs only and positive control is PBMC mixed lymphocyte reaction (MLR) from different donors. Error bars represent mean±SEM from 3-4 donors. Each dot is the mean of two technical replicates. *p<0.05. **(b)** Bar graphs showing percentage of dead PCO from two different lines when co-cultured with CD56+ isolated NK cells from two donors for 4 hours at 37°C at different ratios (1:4, 1:7, 1:15, 1:30 PCO:CD56+). Positive controls are PCO cells treated with tween or kept at 70°C for 3 minutes. **(c)** Bar graphs showing percentage of 7AAD positive dead PCO or ePCO from three different lines when co-cultured with CD56+ isolated NK cells from 3-4 different donors for 4 hours at 37°C at 1:30 organoid:CD56+ ratio. Percentage of dead organoid-only has been subtracted, respectively. Positive controls are PCOs cells treated with tween or kept at 70°C for 3 minutes. Error bars represent mean±SEM from 3-4 donors. *p<0.05.

### HLA DKO ePCOs induce donor-dependent NK cell cytotoxic activity *in vitro*

It is known that lack of HLA-I expression activates NK cells to attack target ‘missing-self’ cells^38,39^. We thus developed a flow cytometric assay to assess the extent of organoid killing by isolated allogeneic CD56+ (NK) cells (**Extended Data Fig. 3b-c**). Overall, parental unedited PCOs were increasingly killed by higher PCO:CD56+ ratios (1:30). However, NK cytotoxicity appeared to be dependent on the donor, potentially influenced by the level of HLA mismatch or NK cell education between NK and PCOs (**Fig. 3b**). NK cells isolated from allogeneic PBMCs induced significantly increased killing of DKO ePCOs in only 1/3 donor-recipient combinations compared to parental unedited PCOs, when co-cultured at ePCO:CD56+ ratio of 1:30. Remarkably, cytotoxicity of NK cells isolated from different donors against the same ePCOs showed a considerably vast heterogeneity (**Fig. 3c**).

### HLA DKO ePCOs show intact graft survival and induce significantly reduced local immune infiltration in humanized mice

We next examined the *in vivo* immune response to PCOs using a humanized mouse model. Fragmented PCOs and ePCOs were injected under the right and left kidney capsules respectively of immunodeficient MHC-I/II dKO NSG mice. At 2 weeks post-injection to allow time for engraftment, animals were humanized by intraperitoneal injection of allogeneic PBMCs obtained from one of two different (allogeneic) human donors (**Supplementary Table 9**). Additionally, one group of mice were injected with Matrigel-only, thus serving as control for the effect of surgery and possible immune response to the Matrigel matrix. 5 weeks post-injection of organoids, animals were culled and their spleens were recovered to assess the human immune cell composition (**Fig. 4a**). Flow cytometry analysis demonstrated increasing levels of human CD45-positive cells in weekly peripheral blood samples, and 50-70% of human CD45-positive cells in the spleen at endpoint, confirming successful humanization (**Fig. 4b, Extended Data Fig. 4a**). Flow cytometry analysis revealed expected reconstitution with predominantly lymphoid cells (CD4, CD8 and B cells) with scarcity of myeloid cells. Both PBMC donors displayed some degree of variability in the immune cell proportions per individual mouse but overall similar mean levels (**Fig. 4c**). Importantly, because each animal was engrafted with both parental PCOs and DKO ePCOs (each line in one kidney of same animal), the experimental design enabled variance in humanization to be controlled for within each animal.

**FIG. 4.**
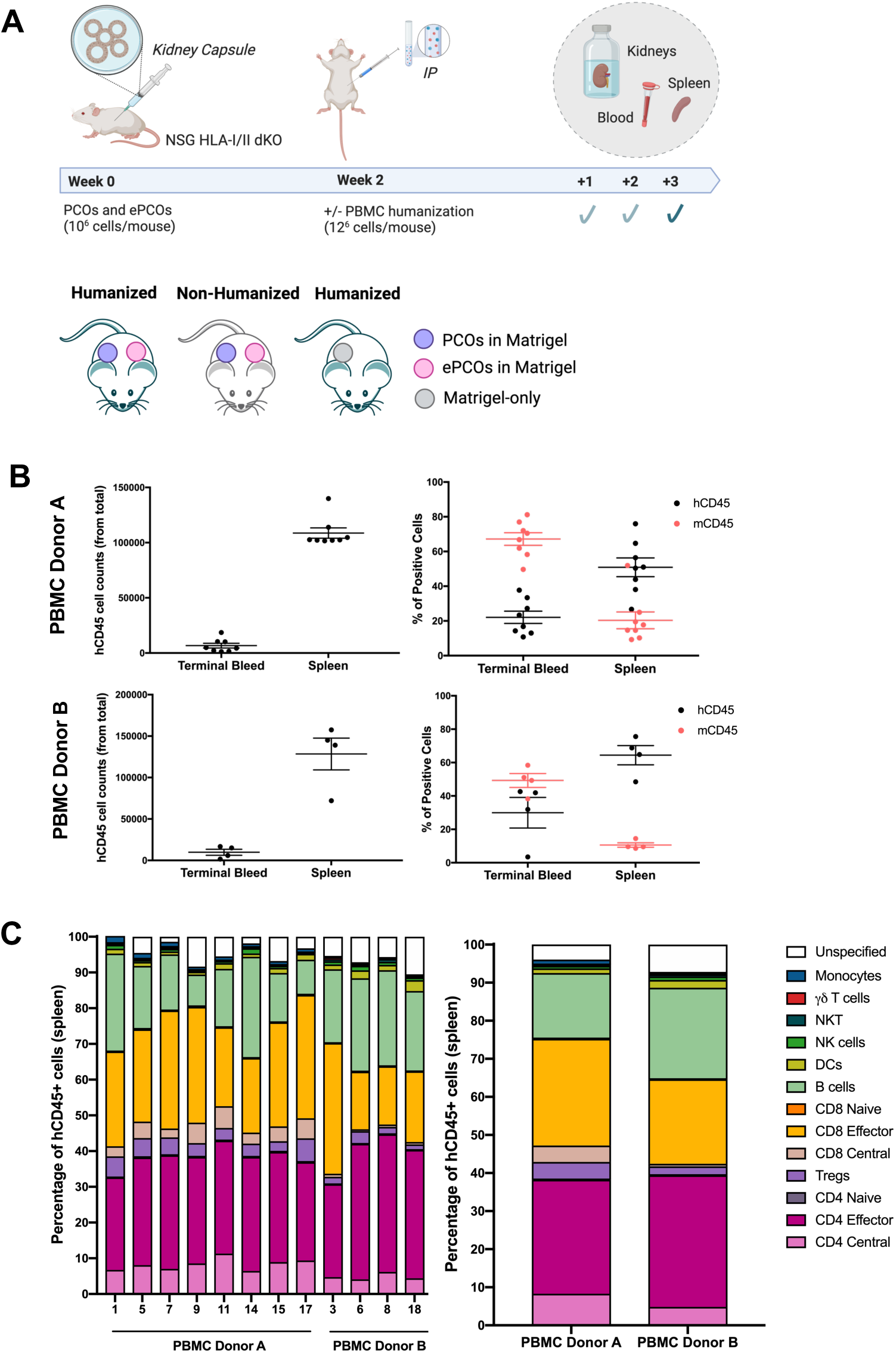
Immune profile of humanized mice transplanted with PCOs and ePCOs. **(a)** Schematics of the PCO injection and subsequent humanization with PBMCs after 2 weeks (n=5 per group). Animals were culled and samples (kidneys, spleens and blood) collected after 3 weeks of humanization (upper panel). Schematic representation of the groups included in the study: PCOs were injected in Matrigel under the right kidney capsule and ePCOs were injected in Matrigel under the collateral left kidney capsule of the same mouse. Matrigel alone was injected under the right kidney capsule of another group of mice to control for Matrigel-driven immune infiltration. Humanization was performed using two different allogeneic human PBMC donors in separate experiments. A non-humanized group was added as control (lower panel). **(b)** Graphs showing human CD45 cell counts (from total) in terminal bleed and spleen at endpoint for the two PBMC donor groups (PBMC Donor A, top; PBMC Donor B, bottom) (left). Graphs showing percentage of positive mouse and human CD45 cells (from live) in terminal bleed and spleen at endpoint (week 5) for the two PBMC donor groups (right). **(c)** Stacked bar graph representing the frequency of immune subsets from human CD45 positive cells in the spleen for each individual mouse and as average from the two PBMC donor groups (right). Error bars represent mean±SEM of 4-8 mice per group.

Assessment of local immune infiltration to the graft site was performed by haematoxylin and eosin staining, human CD45 immunohistochemistry and immunofluorescence staining with specific human immune and cholangiocyte markers (CD45, CD3 and KRT7). Both PCO and ePCO grafts in non-humanized control mice displayed characteristic morphology and surface marker expression (**Fig. 5a**). However, when transplanted into humanized mice, clear differences in PCO and ePCO morphology were apparent in 4/5 animals, with parental unedited PCO grafts displaying an apoptotic-like poor shape, while DKO ePCO grafts maintained the normal morphology observed in non-humanized control mice (**Fig. 5a**, **Table 2, Extended Data Fig. 5a-b, Supplementary Table 10**). There was significantly higher human CD45-positive (predominantly CD3-rich) cell infiltration into the graft region in humanized mice transplanted with parental unedited PCOs compared to DKO ePCOs (**Fig. 5a, Extended Data Fig. 5b**). In some cases, parental PCO grafted organoids had not survived and were not found in the sections (3/8 animals in total across both PBMC donors), while ePCOs survived in all (8/8) animals (**Table 2, Supplementary Table 10**). To quantify immune infiltration, the entire engrafted areas were scanned, hCD45-positive cells were segmented and counted; hCD45-positive stained areas were also quantified with a custom-made pipeline. Importantly, an average quantification of ‘background’ hCD45+ immune infiltration in a group of Matrigel-only control kidneys was used to quantify non-PCO-driven immune infiltration (induced by the surgical procedure or Matrigel itself). The total organoid-targeted immune infiltration was significantly decreased in kidneys that received ePCOs compared to parental PCOs (**Fig. 5b, Extended Data Fig. 5c, Extended Data Fig. 6a-d**). Remarkably, unlike parental PCOs, DKO ePCOs induced an immune infiltration of similar magnitude to that seen in Matrigel-only control kidneys, suggesting effective evasion of the immune response in this humanized mouse model.

**FIG. 5.**
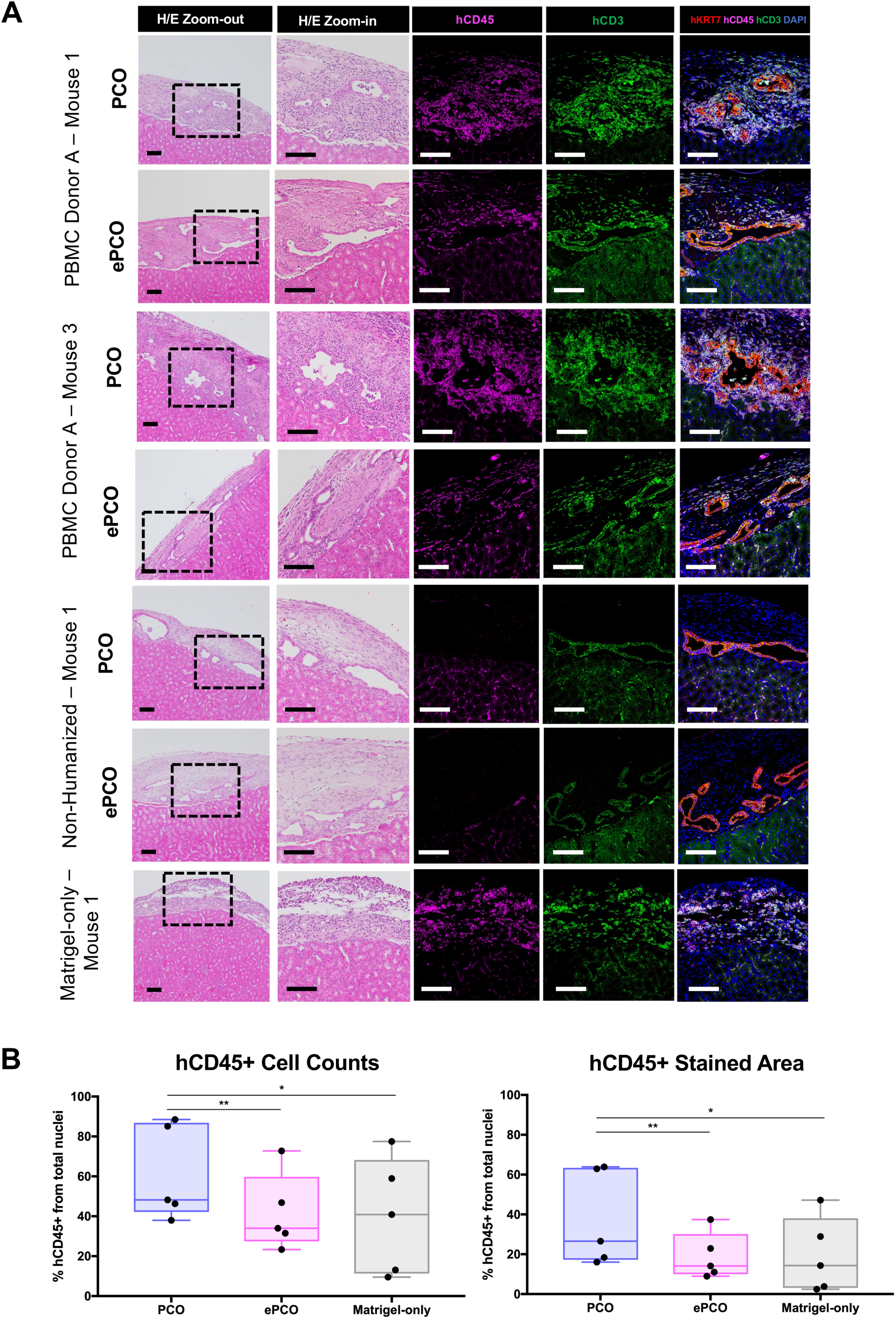
Immune infiltration and graft survival of humanized mice transplanted with PCOs and ePCOs. **(a)** Haematoxylin/Eosin and immunofluorescence staining of PCOs and ePCOs transplanted under kidney capsule showing specific human CD45, CD3 and KRT-7 in PBMC Donor A humanized mice. Non-humanized (positive control for PCO and ePCO engraftment) and Matrigel-only (control for background immune infiltration induced by surgical procedure and injection of Matrigel) are also shown for reference. Scale-bars: 100 µm. **(b)** Percentage of hCD45 positive cells infiltrating into the graft site (left) and percentage of CD45 positive stained area (right) quantified in PCO and ePCO grafted areas of mice humanized with PBMC Donor A. Error bars represent mean±SEM of multiple quantified areas per mouse from 5 mouse per group. *p<0.05; **p<0.01.

**TABLE 2.** PCO and ePCO engraftment in humanized mice with Donor A. Table showing level of PCO and ePCO engraftment (very good, good, poor, not found) in the different mice (M1, M2, M3, M4, M5) humanized with PBMCs from Donor A.

| **Organoid Engraftment** | **PCO** | **ePCO** |
| --- | --- | --- |
| **Non-Hum M1** | Very good | Very good |
| **Non-Hum M2** | Very good | Very good |
| **PBMC Donor A M1** | Poor | Very good |
| **PBMC Donor A M2** | Poor | Good |
| **PBMC Donor A M3** | Poor | Very good |
| **PBMC Donor A M4** | Good | Very good |
| **PBMC Donor A M5** | Not found | Very good |

### ePCO evade T cell mediated cytotoxic immune response, cell graft stress and apoptosis

To understand in more depth the immune response to DKO ePCOs compared to parental PCOs, we carried out spatial transcriptomics specifically for CD45+ (human immune) and PanCK+ (e/PCO) compartments using the NanoString GeoMx DSP platform. Deconvolution analysis revealed that parental PCOs grafted kidneys had at least two-fold more abundant immune infiltration subsets than ePCOs, mostly including central, naïve and effector CD4 and CD8 T cells (**Fig. 6a-b, Extended Data Fig. 7a-b**). Additionally, pathway enrichment analysis of CD45+ (immune) cells showed that human immune cells in ePCOs grafts have downregulated pathways related to T cell activation and cytotoxicity driven by genes coding for perforin (PRF1), IL7 receptor (IL7R) or cathepsin C (CTSC); as well as down regulation of B cell activation and proliferation pathways regulated by genes coding for PTPRC (protein tyrosine phosphatase receptor type C), IKZF3 transcription factor or CD74 receptor (**Fig. 6c, Extended Data Fig. 7c-d, Supplementary Table 11**). Consistent with this immune downregulation, panCK+ (cholangiocyte) compartment of the ePCO grafted animals showed downregulation of pathways involved in cytotoxic cell killing and NF-kappaB-mediated stress response (**Fig. 6d, Extended Data Fig. 7e, Supplementary Table 11**), as also evidenced by the respective graft morphologies described above. Interestingly, KEGG pathway view enrichment results revealed that parental PCO killing mechanism may be driven by extrinsic/receptor-mediated apoptosis pathways (shown by downregulation in ePCO of caspases 6 and 8), while survival of ePCOs may be driven by up-regulation of Bcl-XL and downregulation of pro-apoptotic genes such as p53 (**Fig. 6e**).

**FIG. 6.**
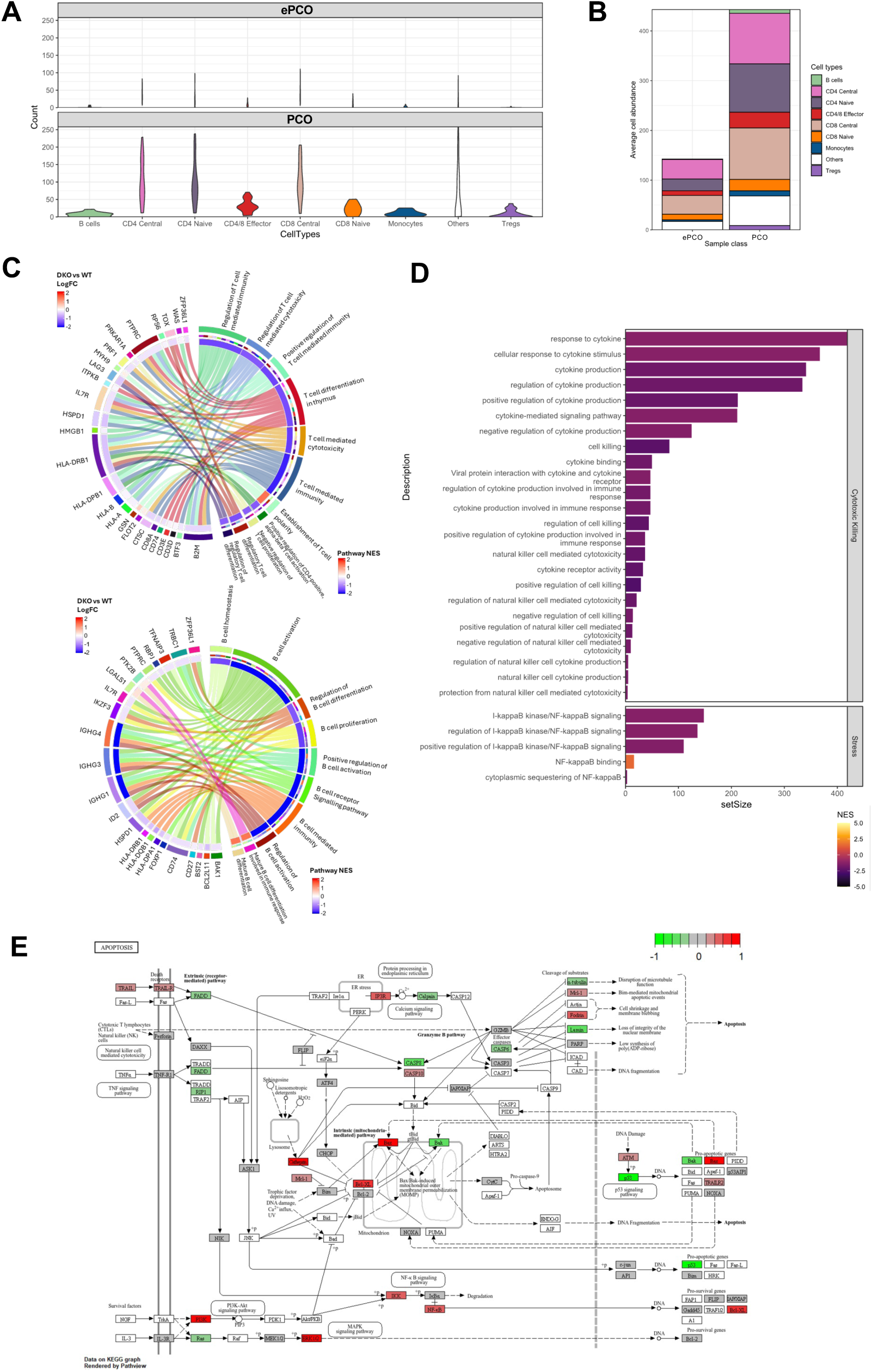
Immune subsets and pathways enriched in CD45+ and PanCK+ compartments upon e/PCO engraftment *in vivo.* **(a)** Violin plot and **(b)** stacked bar chart summarising the average abundance of different immune cell types in DKO ePCO and WT PCO. Matrix (SafeTME) was extracted from doi: 10.1038/s41467-022-28020-5. **(c)** Circos plot showing the GSEA results of ePCO vs PCO CD45+ ROIs for T-cell (left) and B-cell (right) related immune pathway activities and their respective foldchange of pathway-associated genes in CD45+ ePCO vs PCO ROIs. **(d)** Bar chart summarising the GSEA results of ePCO vs PCO PanCK+ cells in apoptosis, cytokine-mediated killing, and stress related pathways. **(e)** KEGG Pathview graph of apoptosis comparing ePCO and PCO PanCK+ ROIs transcriptomics foldchange. Nodes in white or grey indicate values not captured in transcriptomics.

## DISCUSSION

To our knowledge, this work is the first to generate engineered expandable human organoids derived from primary tissue to abolish expression of HLA-I and II molecules aiming to evade immune response. We selected PCOs as our exemplar organoid cellular therapy since they have previously been shown to have potential utility as a cellular therapy^23–25^. In this study, we show that using CRISPR-Cas9 we can eliminate HLA-I and II expression on DKO human PCOs (ePCOs) with minimal off-target mutations, and crucially, without overlapping with known cancer driver genes. Notably, our findings indicate that culturing and passaging of organoids contribute more significantly to mutations than CRISPR off-target edits. However, aided with highly sensitive sequencing methods capable of detecting low-frequency polyclonal mutations, used for the first time in this context, we found no evidence of cancer-driver mutations in the cultures. ePCOs retained morphology, marker expression and functionality of parental organoid cells. Upon *in vitro* co-culture with human immune cells, ePCOs exhibited significantly reduced reactivity, and a donor-dependent NK cytotoxic activity. In the humanized mouse model, ePCOs showed highly preserved graft survival and significantly reduced local immune infiltration compared to parental cells. ePCOs also evaded T cell mediated cytotoxic immune responses, and downregulated cell graft stress and extrinsic apoptotic pathways.

There are several other strategies for reduction or avoidance of immune destruction of cellular therapies. These include the use of an autologous cell source, use of immunosuppressive drugs, generation of HLA-matched donor-recipient banks, and encapsulation of the cells in a barrier material to shield from immune recognition and attack^10^. Genetic engineering allows the development of a limited number of ‘universal’ low-immunogenic engineered organoid cells lines, thus representing important potential economic, logistic and clinical advantages compared to both autologous and fully-immunogenic HLA-typed allogenic cells. Particularly for PCOs, since injectable cholangiocytes must be transplanted in their natural anatomical niche for therapeutic efficacy, their encapsulation in a barrier material is also not compatible with their therapeutic use. Nonetheless, the specific immunomodulatory approach should be chosen based on the inherent properties of the cells, the nature of disease and the organ the cell therapy is aiming to repair.

Gene editing of primary cell-derived organoids presents multiple challenges when compared to engineering pluripotent stem cells. These include determining the right density necessary for single cells derived from organoids to reform as 3D organoids in multiple instances while permitting efficient electroporation and editing conditions (e.g. program, concentration, ratios, guides). In line with few other studies^40–49^, here we show that this process is complex but if optimised can be efficient and successful. We also present *RFX5* as a novel and efficient target for the editing of HLA-II genes and as alternative to *CIITA* previously reported for editing of stem cell-derived cells^14^. Two previous studies described the generation of immune-evasive human organoids: one using overexpression of PD-L1 in iPS cell-derived islet-like organoids^50^, and the other by generating HLA-DKO (*B2M* and *CIITA* edited) with CD47 overexpression in human non-proliferative primary pancreatic islets formed as aggregates of cells^51^. Unlike previous reports, the ePCOs we generated in this study are human primary-derived cholangiocyte cells with comparable 3D morphology and function to primary tissue, and with expandable potential.

Comprehensive characterisation of the off-target effects resulting from any CRISPR-Cas9 editing process is essential to confirm that important genes and cellular programs are not altered. If significant off-target effects are discovered, the affected organoid line should be excluded from further development towards the clinic. Here we explored off-target effects with whole-genome sequencing. Our data suggest that ePCO lines display fewer than 5 mutations in predicted off-target sites and between 5-17 outside predicted off-target sites genome-wide, but without evident impact on functionality or overlap with cancer genes. However, one PCO line acquired an abnormality in chromosome 7 that was not present in the parental PCO line. This may be a spontaneous consequence of cell culture or a pre-existing somatic mutation that expanded *in vitro*, as several studies have previously reported this type of abnormality in cell lines^18,27,28^. Cell lines harbouring such abnormalities should not progress further as a cell therapy, highlighting the need for routine sequencing-based safety assessments in these therapies.

Interestingly, our analyses confirm that time in culture impacts the total mutational burden, in line with previous studies revealing a high mutation rate *in vitro*, associated with oxidative damage in normal oxygen concentration^35–37^. This suggests that total time in culture should be minimised as far as possible, and that lower oxygen concentrations may be important to reduce the mutation rate in culture. Unexpectedly, we noted that ePCOs could present a lower mutation burden than unedited PCOs, a possible explanation being that the cell sorting process can fortuitously select cell clones with a lower mutational burden. Overall, our data suggest that culturing and passaging is a more important source of mutation than CRISPR off-target edits. Reassuringly, targeted NanoSeq analysis of a panel of 257 cancer genes did not detect convincing evidence of cancer-driver mutations in our PCO lines (although this analysis is limited to point mutations and indels).

Our results demonstrate that the majority of mutations acquired by polyclonal organoids during culture cannot be quantified using standard sequencing methods, and highlight the value of duplex sequencing methods including NanoSeq for evaluating mutagenesis in e/PCOs and other organoid systems. These newly developed DNA sequencing technologies offer a high-resolution way of monitoring not only the acquisition of new mutations, but also the clonal selection of cancer-associated mutations present at low allele frequencies. These methods may be critical for future safety studies of cellular products aimed at clinical application.

*In vitro* co-cultures with PBMCs confirmed a significant reduction in immune activation by ePCOs, although there was some activation compared to negative controls (PBMCs alone and organoids alone). This residual immune activation could be driven by reactivity towards dead cells or minor antigens whose surface expression is independent of B2M. NK cell co-cultures showed that ePCOs induced increased cytotoxic activity as expected, although the effect was NK donor-dependent, most likely due to the presence of activating and/or inhibitory NK ligands specific to each donor. It is noteworthy that to demonstrate reproducible NK cytotoxicity, a very high (perhaps unphysiological) ePCO:CD56+ ratio of 1:30 was used. Nonetheless, the fact that ePCOs can induce NK cell cytotoxic activity *in vitro* must be taken into consideration during clinical translation. Possible mitigation strategies could include selection of ePCOs lines with less reactivity to NK cells, to increase dosage of cells administered to account for NK-mediated killing, or to use low doses of immunosuppression that could control the NK-mediated response. A further option would be additional rounds of genetic editing (e.g. knock-in CD47, HLA-E, HLA-G or PD-L1 as previous studies suggested^52–55^). However, the benefits and necessity of additional genetic editing must be balanced against potential safety (mutations acquired in culture) and functional (effect of repeated single cell dissociation) implications for each cellular therapy^18,19^.

We transplanted PCOs and ePCOs under the kidney capsule in humanized mice rather than into the liver. We chose the kidney capsule because it is a highly vascular niche^56,57^ that supports the survival of PCOs, enables localization of the graft at the experimental endpoint, and represents a well-characterized niche for assessment of immunogenicity. Moreover, it enabled PCOs transplanted under one kidney to serve as controls for ePCOs transplanted under the contra-lateral kidney in the same animal. It is possible, however, that the immune response to PCOs may differ qualitatively or quantitatively from our findings when transplanted into the liver in patients. Overall, our humanised mouse model data show that DKO ePCO grafts display a healthier morphology and a higher survival rate than parental controls, which induced more vigorous infiltration with CD45 cells and therefore ongoing immune rejection, in line with other reports with edited pluripotent stem cell-derivatives^58,59^. From the spatial transcriptomics data it is interesting to note the enhanced infiltration of CD45+ delineated histologically corresponded to increased inferred abundance of T cell subsets enriched for cytotoxic activity in the CD45+ compartment of parental PCO grafted areas; and the respective PanCK+ response relating to cell killing, stress and suggesting upregulation of extrinsic apoptosis pathway in response to immune cell interaction. Downregulation of humoral responses was also evident, highlighting how both arms of the adaptive immune response are impacted by HLA-I/II deletion. This indicates that is not simply due to reduced HLA-directed antibody responses, and likely to reflects dampening of tertiary lymphoid structure formation. Although altogether this may seem a reasonable HLA-driven rejection response which can be noticeably evaded by abolishing expression of HLA genes, it is remarkable that human allogeneic response can be captured and neatly described in the humanized mouse model. Future studies utilizing other experimental *in vivo* models that recapitulate the myeloid compartment would be of interest to examine its contribution to ePCO alloimmune response.

Taken together, this work provides first instance of successful HLA-editing in expandable human primary cell-derived organoids, in addition to a comprehensive evaluation of off-target editing and *in vitro* mutagenesis using novel technologies, and of immunogenicity using relevant *in vitro* and *in vivo* experimental models. We believe this study could serve as a guideline for future efforts focused on engineering immune evasive cell-based products for regenerative medicine applications.

## ACKNOWLEDGEMENTS

We are grateful to the donors and their families and the Cambridge Biorepository for Translational Research (CBTM) for the previous gifts of tissue donation. Figures were created in GraphPad Prism and **Fig. 4a** drawings were created with BioRender.com.

S.P.-R. and K.S.-P. are supported by awards from Medical Research Council (MRC) UK Regenerative Medicine Platform (MR/S020934/1) and MRC Confidence in Concept (G116517). I.M. is funded by Cancer Research UK (C57387/A21777), the Dr Josef Steiner Cancer Research Foundation and the Wellcome Trust. M.J.B. and G.K. are funded by the British Skin Foundation (034/YI/22) and the Chinese Academy of Medical Sciences (CAMS) Innovation Fund for Medical Science (CIFMS), China (2018-I2M-2-002).

## AUTHOR CONTRIBUTIONS

S.P.-R. conceived and designed the study, performed experiments, acquired, interpreted, and analysed the data, drafted and edited the manuscript. A.B.-O. designed and performed bioinformatic mutation analyses, and edited the manuscript. W.L. performed bioinformatic spatial transcriptomic analysis. G.K. performed sample preparation for spatial transcriptomic analysis. J.W. performed and analysed proteomics experiments and data. J.J. and C.B. performed slide scanning and hCD45 quantification. D.T. contributed to *in vivo* experiments. A.R.J.L. contributed to sample preparation for sequencing and to targeted nanoseq. K.M. provided primary tissue. P.L. provided funding, designed and interpreted proteomic experiments. M.J.B. helped designing spatial transcriptomic experiment, interpreted data and edited the manuscript. I.M. provided funding, designed off-target experiments, interpreted data and edited the manuscript. K.S.-P. Conceived and designed the study, performed animal experiments, interpreted data, wrote and edited the manuscript. All authors approved the manuscript.

## EXTENDED DATA FIGURES

**EXTENDED DATA FIG. 1.**
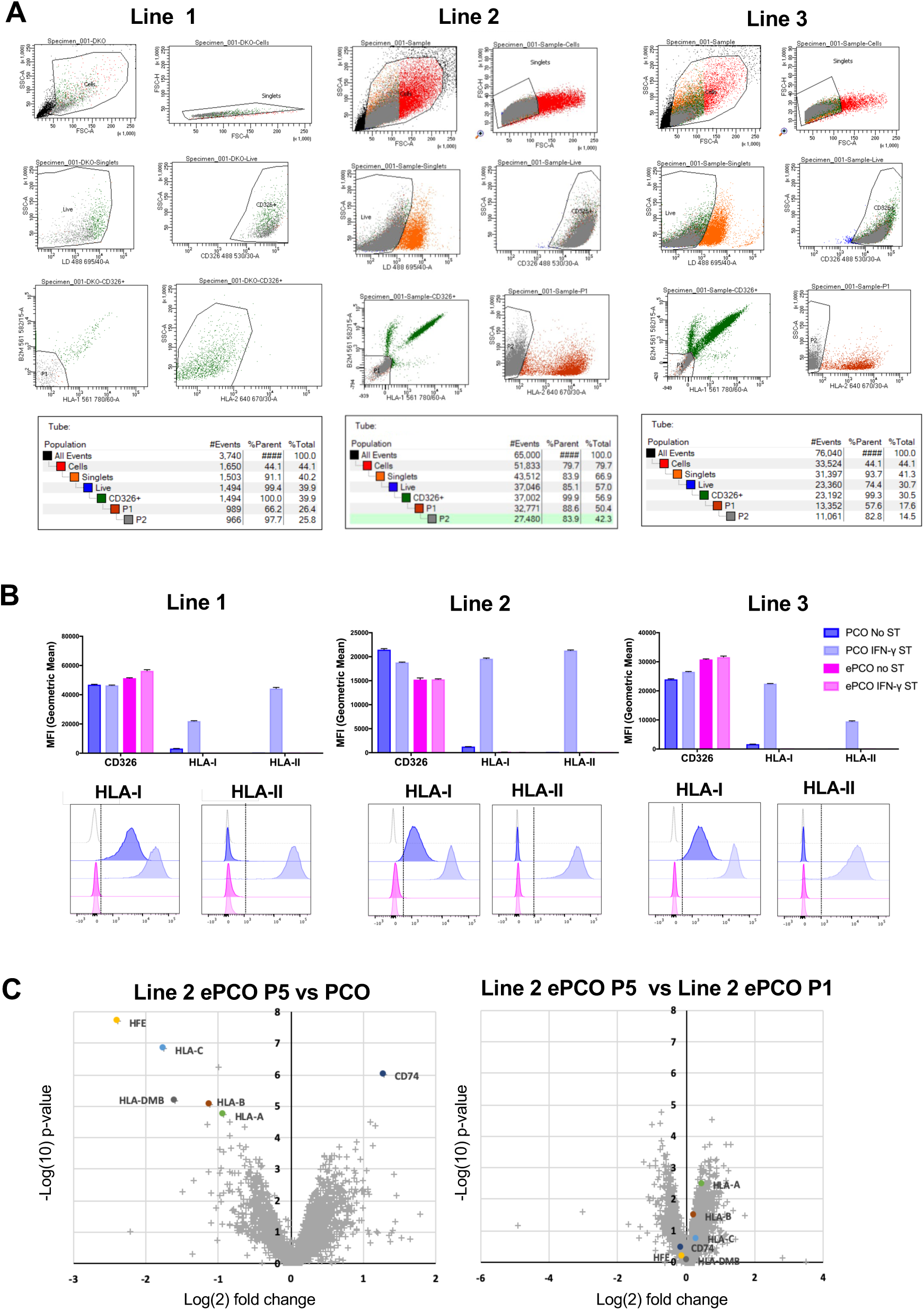
Generation and characterisation of ePCO HLA-I/II DKO lines. **(a)** Gating strategy and respective population fractions used at cell sorting to generate the three lines of engineered ePCOs. **(b)** Bar graphs showing mean intensity fluorescence (MFI, top) and respective flow cytometry charts (bottom) of CD326, HLA-I and HLA-II for PCO and ePCO organoids cultured with and without IFN-ɣ stimulation (2 days, 100ng/mL) for the three lines. Fluorescence minus one (FMO) controls are used for gating. Error bars represent mean±SEM from three independent experiments. **(c)** Plots showing differentially expressed surface proteins in Line 2 ePCO passage 5 (P5) compared to PCO, and Line 2 ePCO passage 5 (P5) compared to Line 2 ePCO passage 1 (P1). Cut-off q-value of <=0.01 and fold change cut off of 2-fold.

**EXTENDED DATA FIG. 2.**
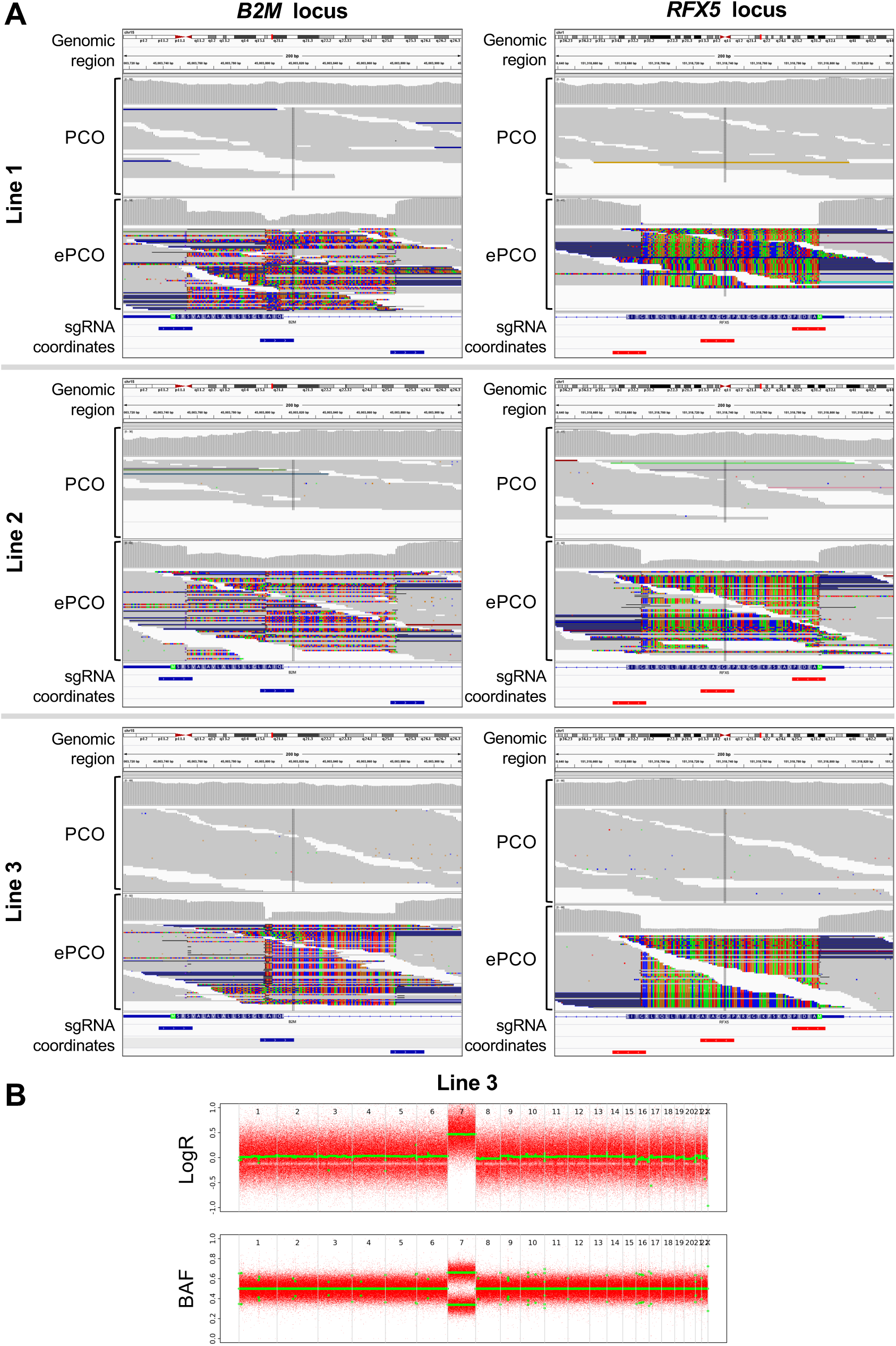
Evidence of CRISPR DKO edits and copy number alterations from WGS. **(a)** Integrative Genomics Viewer (IGV) read alignment images showing CRISPR KO events in *B2M* (left; chr15:45,003,717-45,003,916) and *RFX5* (chr1:151,318,639-151,318,838) for PCO and ePCO samples from each engineered line. Soft-clipped portions of sequence reads (indicating CRISPR-induced deletions) are coloured according to base type. Sequencing depth is represented in grey above each read alignment. Gene annotation and single guide RNA (sgRNA) coordinates are shown at the bottom of each panel. No single sequence read spans the entire region between the sgRNAs without soft-clipping, suggesting that all sequenced cells carry a deletion in the target region. **(b)** Values of log-transformed sequencing depth ratio (logR) and B-allele frequency (BAF) along the genome of the ePCO sample from Line 3, indicating whole-chromosome gain (trisomy) of chromosome 7.

**EXTENDED DATA FIG. 3.**
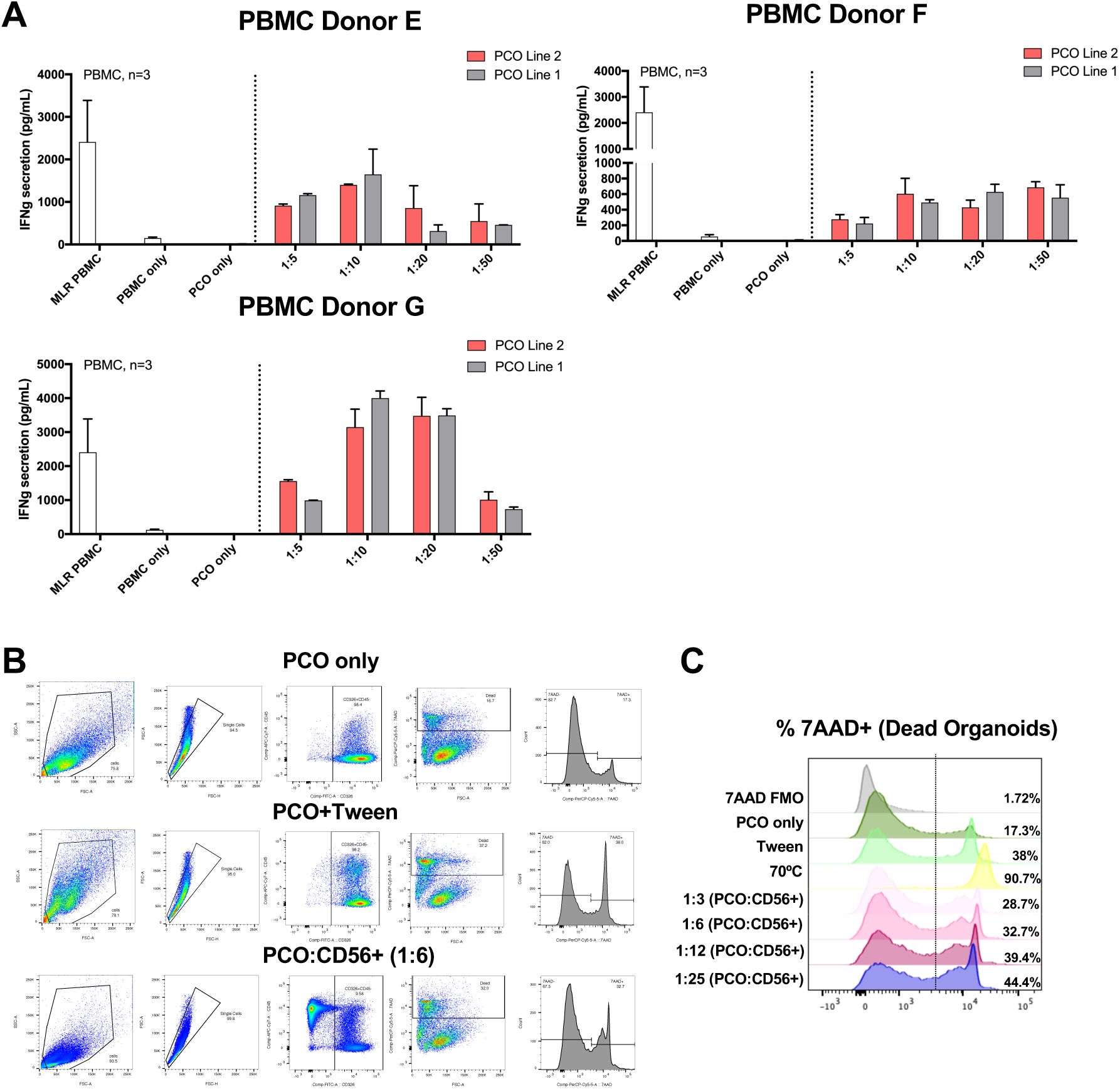
Optimisation of the co-culture assays with human immune cells. **(a)** Bar graphs showing concentration of IFN-ɣ secretion by PBMCs when co-cultured at different PCO:PBMC ratios (1:5, 1:10, 1:20, 1:50) with IFN-ɣ pre-stimulated PCOs for 5 days. Negative controls are PCOs only and PBMCs only and positive control is PBMC MLR from different donors. Error bars represent mean±SEM from technical triplicates. **(b)** Gating strategy used to determine percentage of dead PCOs. As reference, PCO only, PCO cells treated with tween (positive control) and PCOs mixed with CD56+ isolated NK cells (1:6 PCO:CD56+ ratio). **(c)** Representative flow charts showing percentage of dead PCOs at different PCO:CD56+ ratios. Positive controls are PCO cells treated with tween or kept at 70°C for 3 minutes. Fluorescence minus one (FMO) control is shown as reference.

**EXTENDED DATA FIG. 4.**
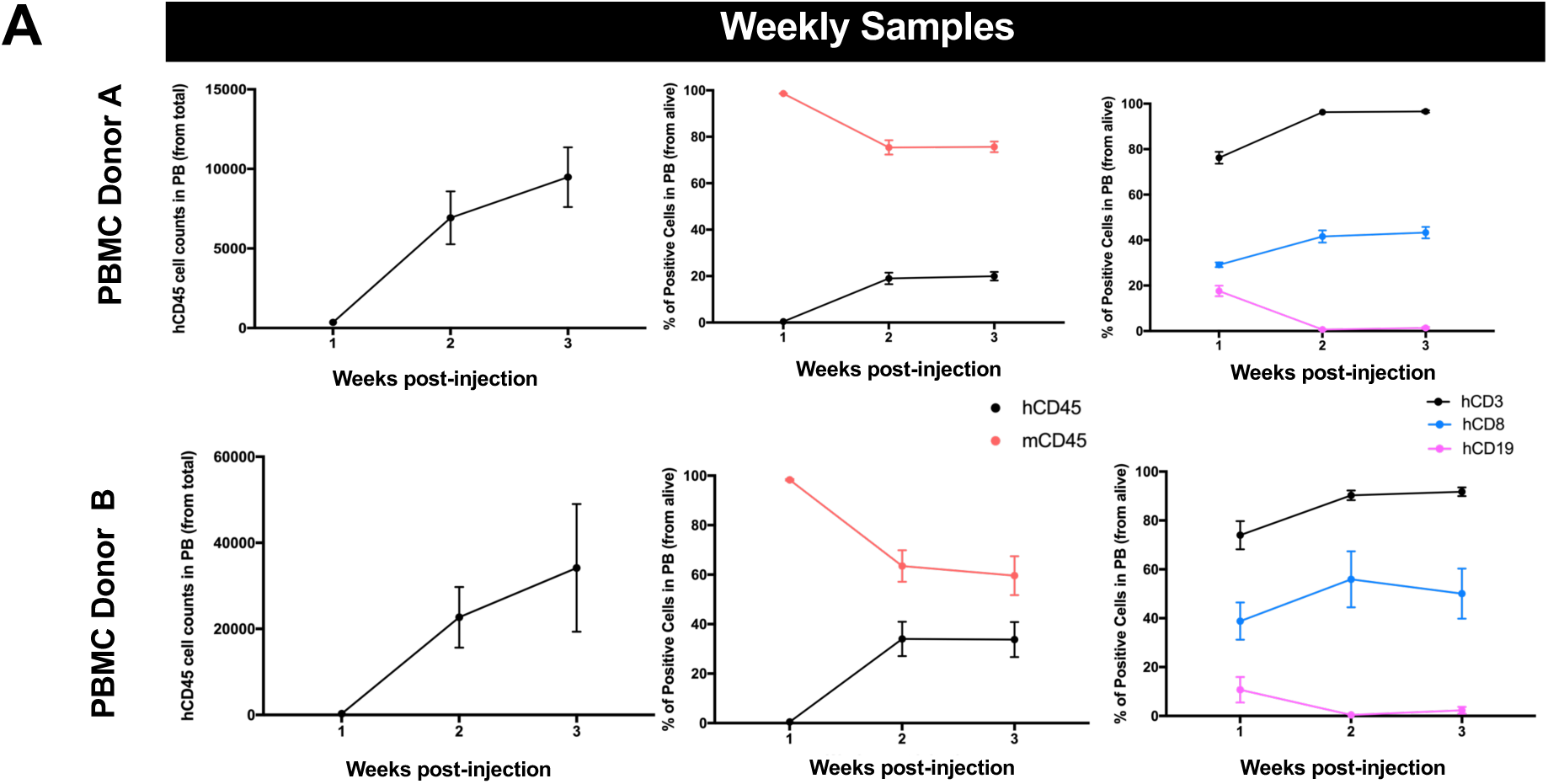
PBMC donor engraftment in NSG HLA-I/II dKO mice. **(a)** Graphs showing percentage of positive mouse and human CD45 cells, percentage of human CD3, CD8, CD19 and human CD45 cell counts in weekly in peripheral blood (tail vein bleeds) for the two PBMC donor groups (PBMC Donor A, top; PBMC Donor B, bottom). Error bars represent mean±SEM of 4-8 mice per group.

**EXTENDED DATA FIG. 5.**
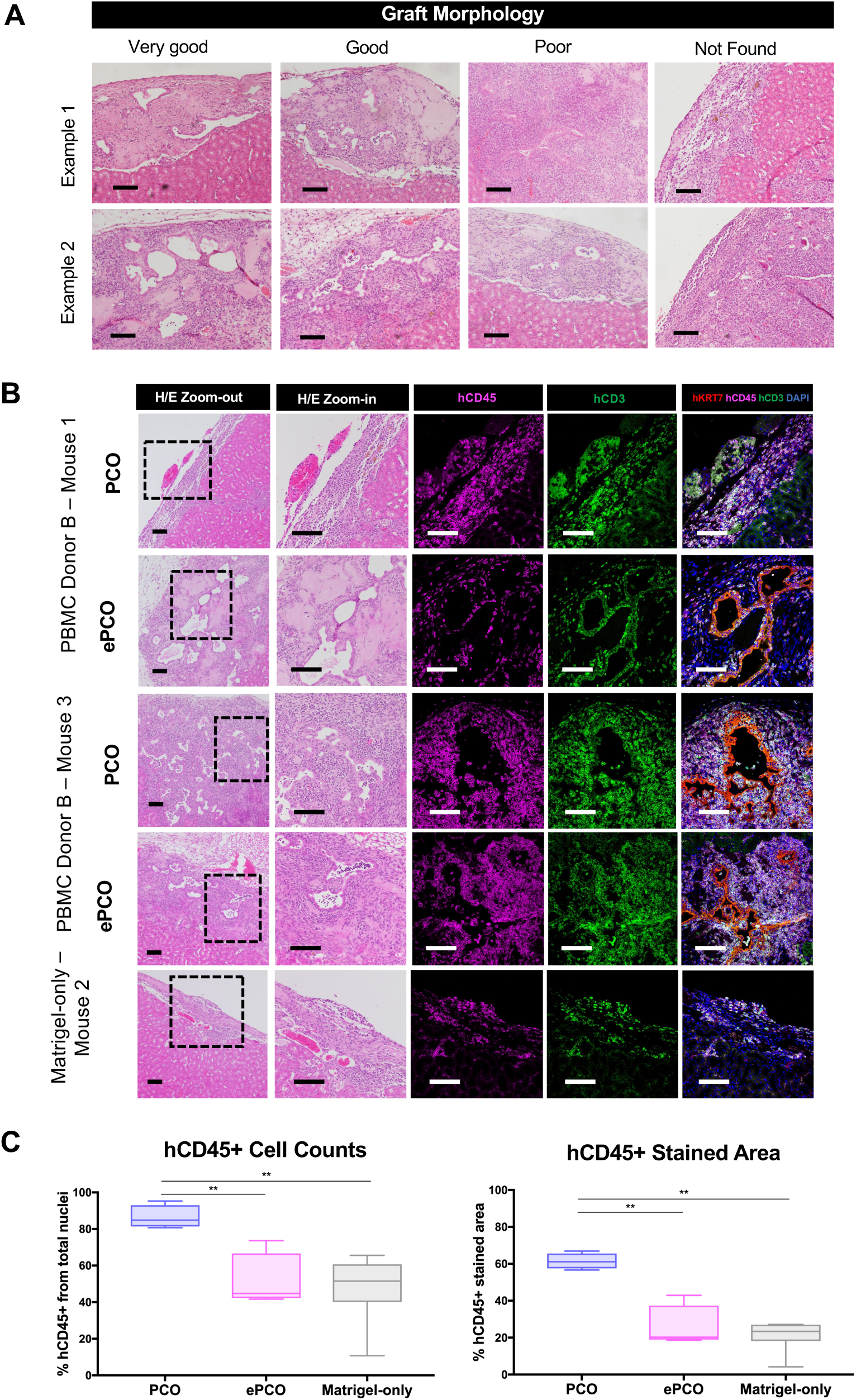
Local immune infiltration and graft survival of humanized mice transplanted with PCOs and ePCOs. **(a)** Haematoxylin/Eosin images representing examples of very good, good, poor and not found PCO engraftments under the kidney capsule of humanized mice. Scale bars: 100 µm. **(b)** Haematoxylin/Eosin and immunofluorescence staining of PCOs and ePCOs transplanted under kidney capsule showing specific human CD45, CD3 and KRT-7 in PBMC Donor B humanized mice. Non-humanized (positive control for PCO and ePCO engraftment) and Matrigel-only (control for background immune infiltration induced by surgical procedure and injection of Matrigel) are also shown for reference. Scale bars: 100 µm. **(c)** Percentage of hCD45 positive cells infiltrating into the graft site (left) and percentage of CD45 positive stained area (right) quantified in PCO and ePCO grafted areas of mice humanized with PBMC Donor B. Error bars represent mean±SEM of multiple quantified areas per mouse from 3 mouse per group. **p<0.01.

**EXENDED DATA FIG. 6.**
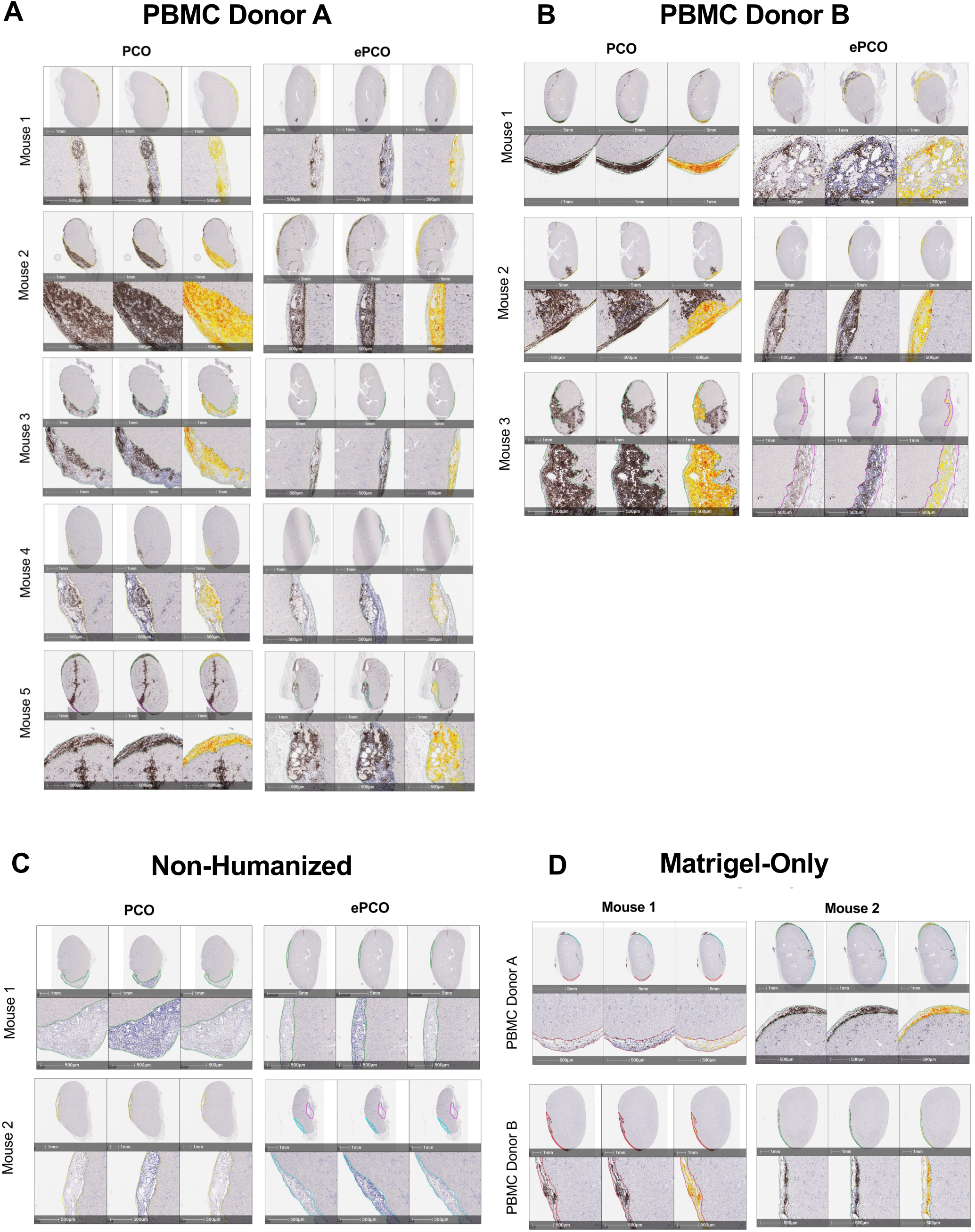
CD45+ immune infiltration in humanized mice with transplanted PCOs and ePCOs. **(a)** Low (upper rows) and high (lower rows) magnification scans and respective custom-made segmentation for CD45 cells (middle) and area quantification (right) showing positive human CD45 staining (brown, left) and hematoxylin staining (nuclei, left) in kidneys from several mice that received PCO (left panels) or ePCO (right panels) for Donor A or Donor B **(b). (c)** Non-humanized and Matrigel-only **(d)** groups are shown for reference. Marked areas in green, magenta and yellow are the ones used for further quantification. Scale-bars: 500µm, 1mm and 5mm.

**EXENDED DATA FIG. 7.**
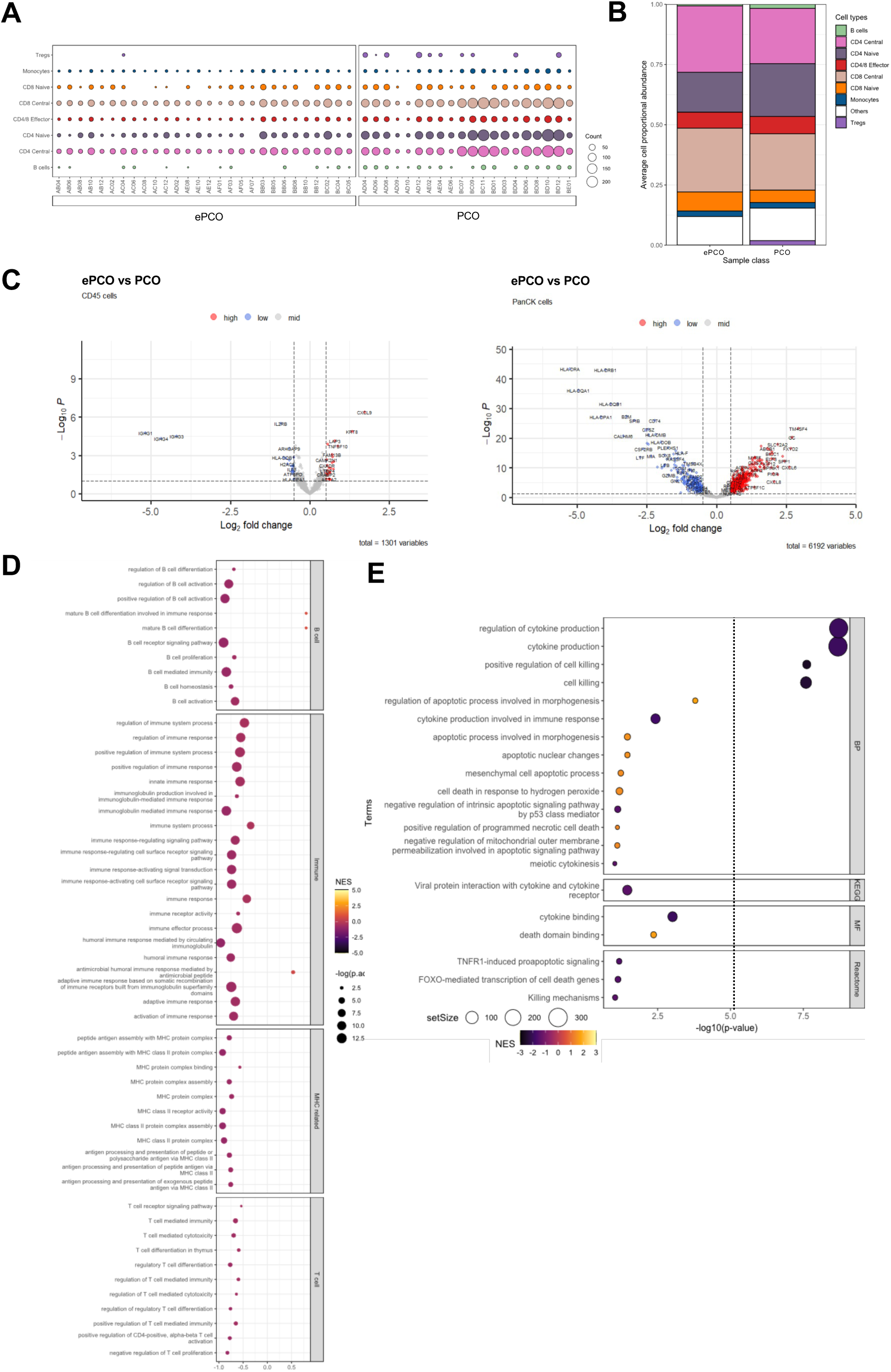
Immune subsets and pathways enriched in CD45+ and PanCK+ compartments upon e/PCO engraftment *in vivo.* **(a)** Dot plot showing the abundance per sample of different immune cell types in DKO ePCO and WT PCO. Matrix (SafeTME) was extracted from doi: 10.1038/s41467-022-28020-5. **(b)** Stacked bar summarising the average cell proportional abundance of different immune cell types in ePCO and PCO. **(c)** Volcano plot showing low, mid and high differentially expressed genes in CD45+ (right) and PanCK+ (left) cells compared to PCO. **(d)** Dot plot summarising the GSEA results of ePCO vs PCO CD45+ ROI in B cell, Immune, MHC-related and T cell related pathways. **(e)** Summary of ePCO vs PCO PanCK+ ROI GSEA biological process (BP), molecular function (MF), KEGG and Reactome results in apoptosis, cytokine-mediated killing, and stress related pathways. Dotted line to the right marks significant p-value (<0.05).

## METHODS

### PCO Derivation and Passage

Cholangiocyte lines were derived from tissue gifted by deceased transplant organ donors under ethical approval (NRES Committee East of England - Cambridge South, Ref 15/EE/0152) and with informed consent from the donor families. After retrieval, extrahepatic bile ducts were collected, cut as flat sheets with sterile scalpel, and washed three times in separate tubes with PBS (Sigma, D8537). In a petri dish with cholangiocyte media (CM, see reference Tysoe et al.^24^), cells from the lumen were scraped until velvet structure was not visible. Tissues were flushed with CM and all media was collected and centrifuged 5 min, 400g. If pellet appeared red, a red cell lysis removal step with ammonium chloride solution (Stem Cell Technologies, 07850) was performed (12min, 4°C). After, cells were washed twice with CM and debris and fibrous tissues pieces were removed with pipette. Plating was performed according to pellet size resuspending cells in 2:3 of Matrigel (Corning, 356237) and 1:3 of complete cholangiocyte media (CCM) consisting of hDKK 100ng/mL (R&D Systems, 5439-DK-01M/CF); hEGF 50ng/mL (R&D Systems, 236-EG-01M); hR-Spondin-1 500ng/mL (R&D Systems, 4645-RS/CF) and Rock Inhibitor-Y27632 10mM (Selleckchem, S1049). 50µL - domes were plated in 24-well or 6-well plates kept at 37°C for 3min and subsequently flipped up-side-down at 37°C for 15 min. CCM was then added to the well at a final volume of 1mL (24-well) or 3mL (6-well), and cultures were maintained in CCM without Rock Inhibitor-Y27632 in a 5% CO_2_ incubator. Typical PCO morphology appeared after 2-3 days and organoids were ready to passage after 7-10 days.

Established PCO lines were passaged mechanically at 1:3-1:5 ratio every 3-5 days as follows: organoids were retrieved from domes in a collection tube using Cell Recovery Solution (Corning, 354253). Pooled domes in the solution were kept on ice for 15 min. Tubes were centrifuged for 5 min, 400g and washed with 1mL of CM. Pellet size were assessed and divided in tubes for subsequent passage. PCO pellets were resuspended with 1:3 of CCM and mechanically fragmented with a p200 (40-60 times) until middle-size homogenous fragments were observed under the microscope. Then, 2:3 of Matrigel was added to the fragmented PCOs per tube, resuspended well, and 50µL-domes were plated in 24-well or 6-well plates. Plated domes were kept at 37°C for 3 min and subsequently flipped up-side-down at 37°C for 15 min. CCM media was then added to the well to a final volume of 1mL (24-well) or 3mL (6-well). Cultures were fed three times a week with CCM without Rock Inhibitor-Y27632 in a 5% CO_2_ incubator.

### PCO CRISPR-Cas9 Editing

PCOs were retrieved from domes and pellets were resuspended with 1mL of Accutase (Sigma, A6964), DNAse-I (NEB, Sigma, 4536282001) and Rock Inhibitor-Y27632 10mM (Selleckchem, S1049) per tube. Tubes were placed in a water bath at 37°C for 5 min, followed by 10-times gentle resuspension. The single cell suspension was evaluated under microscope until clumps were not seen (if needed, tubes were placed in a water bath 37°C for extra 5 min). 3mL of CM media was added to stop the reaction and cells were centrifuged for 5 min 400g. Cells were filtered with a 40µm cell strainer (FisherBrand, 22363547) and counted with Countess II FL (Life Technologies).

Subsequently, RNPs were pre-complexed with single guide RNA (sgRNA, Synthego Gene Knockout Kit, Synthego) and TrueCut Cas9 Protein v2 (A36498, ThermoFisher Scientific), incubated for 10 min at room temperature (see commercially available guides in **Suppl Table 2**). Complexed RNPs were re-suspended in 100µL of P3 buffer including supplement 1 (P3 Primary Cell 4D-Nucleofector Kit, V4XP-3024, Lonza) and electroporated using 4D-Nucleofector^TM^ System X Unit (Lonza). Finally, 5 × 10^4^-3 × 10^5^ cells were plated in 50µL domes containing 2:3 of Matrigel (Corning, 356237) and 1:3 of complete cholangiocyte media (see Tysoe et al^24^). Cells were placed at 37 °C and 5% CO_2_.

Organoids start to reform after 3 days after electroporation. Knockout efficiency was further evaluated using flow cytometry for HLA-I and HLA-II molecules (see below). To increase purity, organoids were sorted using flow cytometry cell sorting and gating on the negative population of interest.

For the double knock-out generation a sequential approach was taken, therefore organoids were first electroporated with B2M-RNP complexes, grown and further sorted for B2M-/HLA-I-; and subsequently, they were additionally electroporated with RFX5-RNP complexes, grown and further sorted for both HLA-I-/HLA-II-populations (see below).

### Flow Cytometry and Cell Sorting

PCOs were retrieved from domes as described for passaging and pellets were resuspended with 1mL of Accutase (Sigma, A6964), DNAse-I (NEB, Sigma, 4536282001) and Rock Inhibitor-Y27632 10mM (Selleckchem, S1049) per tube. Tubes were placed in a water bath at 37°C for 5 min, followed by 10-times gentle resuspension. The single cell suspension was evaluated under microscope until clumps were not seen (if needed, tubes were placed in a water bath 37°C for extra 5 min). 3mL of FACS Buffer (2% FBS (Thermo Fisher Scientific, 10082-147) and 1mM EDTA (Thermo Fisher Scientific, 15575-020) diluted in DPBS (Sigma, D8537)) was added to stop the reaction and cells were centrifuged for 5 min 400g. Cells were filtered with a 40µm cell strainer (FisherBrand, 22363547), centrifuged and further stained using the following conjugated antibodies: mouse anti-human HLA-ABC-PeCy7 (1:20, BD Biosciences, 561349, clone [G46-2.6]), HLA-DR, DP, DQ-APC (1:20, Biolegend, 361714, clone [Tü39]) and CD326-FITC (1:20, Biolegend, 324204, clone [9C4]).

Cells were incubated with the conjugated antibodies at 4°C for 30 min and washed twice with FACS Buffer. Fluorescence minus one (FMO) controls were included for each condition to gate negative and positive cells. 7-AAD viability staining solution (1:50, Biolegend, 420404) was added to the cells. Stained cells were analyzed using a FACS Canto II (BD Biosciences). Analysis of the data was carried out using FlowJo v10 software (Tree Star). Biological triplicates were used for every condition. Results are presented as mean±SEM (standard error of the mean).

For sorting, expanded organoids were brought to single cells and incubated with the mentioned conjugated antibodies on ice for 30 min. Fluorescence minus one (FMO) controls were included for each condition to identify and gate negative and positive cells. Stained cells were then sorted using a BD FACS Aria Fusion Cell Sorter (BD Bioscience) using FACSDiva Software v8.0.1. After sorting, 5 × 10^4^-3 × 10^5^ of DKO cells were plated in 50µL domes containing 2:3 of Matrigel (Corning, 356237) and 1:3 of complete cholangiocyte media Rock Inhibitor-Y27632. Cells were placed at 37 °C and 5% CO_2_. Typical PCO morphology appeared after 5-10 days.

### HLA Genotyping

Genomic DNA was extracted from either cell pellets was isolated using QIAmp DNA Blood Mini Kit (Qiagen, S1104) following manufacturer’s instructions; or cell suspensions following manufacturer’s instructions of EZ1&2 DNA Blood 350µl Kit (Qiagen, 951054) and EZ1 Advanced XL instrument (Qiagen). DNA was quantified and purity assessed using NanoDrop One/OneC Microvolume UV-Vis Spectrophotometer (Thermo Fisher Scientific). Multiplex HLA amplification of 11 loci (HLA-A*, B*, C*, DRB1*, DRB3/4/5*, DQA1*, DQB1*, DPA1* and DPB1*) was performed with GenDx NGSgo-MX11-3 kits (GenDx, 7971864). The library was quantified using Qubit dsDNA (broad range) BR Assay Kit (Invitrogen, Q32850) and the final library loading concentration was 15pM. Sequencing was performed using MiSeq instrument (Illumina) and MiSeq Reagent Kit v2 (Illumina, MS-102-2002) with 300 cycle flow cells. Analysis was done using GenDx NGSengine software (version 2.28.1; 3.50 IMGT/HLA release version) complying with European Federation of Immunogenetics (EFI) accreditation.

### Gamma Glutamyl Transferase (GGT) Assay

Cultured organoids (1-3 domes per group per line) were homogenized in 200µL of GGT assay buffer from Gamma Glutamyl Transferase Assay Kit (Colorimetric; abcam, ab241028) and centrifuged at 13,000g for 10 min to remove insoluble material. 10µL of the resulting supernatant was taken to a 96-well plate to proceed with the kit as indicated in manufacturer’s instructions (abcam, ab241028). Optical density was measured at 405nm in a microplate reader (BMG LABTECH 96) after 90 min incubation at 37°C with 90µL of GGT substrate mix. Samples were analysed in triplicates and linear regression with blank correction was applied.

Primary bile duct/gallbladder snap frozen tissue was used as primary material control, in addition to a GGT positive control included in the assay and substrate only control as negative control. Results are presented as mean±SEM (standard error of the mean).

### Alkaline Phosphatase (ALP) Staining

Cultured organoids (1 dome/24-well per staining per group per line) were stained using Alkaline Phosphatase Staining kit (abcam, ab242287) following manufacturer’s instructions. Domes (in matrigel) were fixed for 5 min with fixing solution, washed twice with PBS (Sigma, D8537) containing 0.05% Tween-20 (Sigma, P1379) and stained with AP staining for 45 min, followed by two washings with PBS. Samples without AP staining were used as negative control. Samples were imaged after 1-2 with an Olympus IX81 fluorescence inverted microscope. Post-acquisition analysis was performed using ImageJ v2.0 software.

### PBMC Isolation from Human Donors

Human PBMCs were isolated from leukopaks of healthy donors firstly removing red blood cells by adding ammonium chloride solution (StemCell Technologies, 07850) to the sample at a volume:volume ratio of 4:1 and incubated for 15min on ice. Samples were centrifuged at 500g for 10min at room temperature. An additional wash with 2% FBS in PBS at 200g for 10min at room temperature with the brake off was carried out to remove platelets. Cells were counted with Countess II FL (Life Technologies) and if not used directly, they were frozen in 10% DMSO (Sigma, D8418-250mL) in FCS accordingly (50 million cells/mL per vial).

### Isolation of Human NK cells from Healthy Human Donors

Human PBMCs were isolated as described above and consecutively NK cells were negatively separated with a CD56 isolation kit (Miltenyi Biotec, 130-050-401) using autoMACs Pro Separator (Miltenyi Biotec) with the “depletes” program. Purities were of above 90% for CD56+ CD3-selected cells in the donors used for the experiments (data not shown). Final cell numbers were assessed Countess II FL (Life Technologies) and cells were seeded out at a concentration of 1×10^6^ cells/mL in RMPI medium (Thermo Fisher Scientific, 21875-034) with 10% AB serum (Sigma, H3667) and 100U/mL penicillin-streptomycin (Thermo Fisher Scientific, 15140-122), and activated over night with 1000 U/mL IL-2 (R&D Systems, 202-GMP-050).

### Co-culture of PCOs with PBMCs

PBMCs were first thawed in a water bath 37°C, washed with 50% FBS in RPMI followed by 10% FBS in RPMI wash, and filtered through a 40µm strainer (FisherBrand, 22363547). Cell numbers were assessed by counting with Countess II FL (Life Technologies). 2 days IFN-ɣ pre-stimulated (100 ng/mL, R&D Systems, 285-IF-100) organoids were single celled as described above and plated at a cell density range of 9×10^5^ cells/cm^2^ (1:10, organoid:PBMC) using a 1:1 mix of CCM media and RPMI medium (Thermo Fisher Scientific, 21875-034) with 10% AB serum (Sigma, H3667) and 100U/mL penicillin-streptomycin (Thermo Fisher Scientific, 15140-122). 1.5×10^6^ of isolated PBMCs were plated per well in a 48-well plate on top of the plated single cell organoids. IL-2 (1ng or 100U, BD Biosciences, 554603) and CD28 (1.25μg/mL, Biolegend, 302902 [CD28.2]) molecules were added to the wells. For MLR, PBMCs from the different donors were mixed together 1:1 with the addition of the stimulatory molecules. Co-cultures were maintained for 5 days at 37°C for further analysis.

### Co-culture of PCOs with CD56+ NK cells

NK mediated cytotoxicity was measured by percentage of dead organoids by flow cytometry. Overnight IL-2 activated hNK cells (effector cells) were mixed with 2 days IFN-ɣ pre-stimulated (100 ng/mL, R&D Systems, 285-IF-100) e/PCOs (target cells) brought into single cells (see above) at different target(e/PCO):effector(CD56+) ratios (depending on the experiment; 1:4; 1:7; 1:15; 1:30) in a 96-well plate with RPMI medium (Thermo Fisher Scientific, 21875-034) with 10% AB serum (Sigma, H3667) and 100U/mL penicillin-streptomycin (Thermo Fisher Scientific, 15140-122). Plate was briefly centrifuged at 100g for 10 sec and incubated for 4 hours at 37°C.

After, cells were centrifuged at 300g for 7 min and further stained using the following conjugated antibodies: mouse anti-human CD326-FITC (1:20, Biolegend, 324202 clone [9C4]), CD56-APC (1:20, Biolegend, 318310, clone [HCD56]), CD45-APC/Cy7 (1:20, Biolegend, 304014, clone [HI30]) and CD3-PE/Cy7 (1:20, Biolegend, 300420, clone [UCHT1]). Cells were incubated with the conjugated antibodies at 4°C for 30 min, washed with FACS Buffer and were filtered with a 40µm strainer (FisherBrand, 22363547). Fluorescence minus one (FMO) controls were included for each condition to gate negative and positive cells. Also, e/PCO only and CD56+only cells were added as controls. 7-AAD viability staining solution (1:50, Biolegend, 420404) was added to the cells. Stained cells were analyzed using a FACS Canto II (BD Biosciences). Analysis of the data was carried out using FlowJo v10 software (Tree Star). Results are presented as mean±SEM (standard error of the mean).

### Enzyme-Linked Immunosorbent Assay (ELISA)

Supernatants from organoids and PBMC co-cultures were collected 5 days after the cells were plated. Cytokine secretion levels (IFN-ɣ, TNF-α, IL-6, IL-10, IL-13, IL-1β, IL-2, IL-4, IL-8, IL-12p70) were measured for each condition in duplicates with commercially available 10-plex human Proinflammatory panel (MesoScale Discovery, K15049D-2) in accordance with the instructions of the manufacturers. Samples were diluted 1:2 in assay diluent. The electrochemiluminescence readings were measured using s600 reader (MesoScale Discovery). Results are presented as mean±SEM (standard error of the mean).

### *In vivo* Experiments

Animal experiments were conducted in accordance with UK Home Office and Institutional regulatory requirements (HO PP5753595). Male and female immunodeficient Nod Scid Gamma (NSG) mice aged 6-10 weeks were used in this study.

### PCO Injections and Humanization

An average of 15 domes of PCOs and ePCOs were used for transplantation into each animal. Organoids were retrieved from domes in a collection tube using Cell Recovery Solution (Corning, 354253). Pooled domes in the solution were kept on ice for 15 min. Tubes were centrifuged for 5 min at 400g and washed with 1mL of CM. Organoid pellets were resuspended with CCM and mechanically fragmented with a p200 pipette (40-60 times) until middle-size homogenous fragments were observed under the microscope. All fragments were combined together and well-resuspended with 1:3 of CCM media and 2:3 of Matrigel (Corning, 356237). Cells were transported on ice and loaded into a 50mL syringe (Hamilton 700/1700 Series Microliter/Gastight Syringes, Model: 1705) with a blunt needle (custom-made, FisherScientific, 10698205). 25µL of cell mix was injected under the left or right kidney capsule following a flank incision to deliver the kidney onto the operating field under isoflurane anaesthesia. Matrigel-only was injected under the right kidney capsule as control in some animals. After surgery, animals were administered additional saline and mesh food, and were closely monitored for any adverse effects.

2 weeks after cell injection, animals (n=5 per group) were humanized with 12×10^6^ human PBMCs per mouse. PBMCs were thawed in a water bath at 37°C, washed with 50% FBS in RPMI followed by 10% FBS in RPMI wash, and filtered through a 40μm strainer (FisherBrand, 22363547). Cell numbers were assessed by counting with Countess II FL (Life Technologies). Total amount of cells needed were resuspended in total volume needed of 2% FBS in PBS and kept on ice. 200µL of PBMCs were injected intraperitoneally per mouse with a 29g needle (Beckton Dickinson, 324892). Humanization was evaluated weekly by flow cytometry of tail-vein bleeds collected in heparin-coated tubes (Sarstedt Ltd, 20.1309), followed by red cell lysis removal (StemCell Technologies, 07850) according to manufacturer’s instructions and using the following conjugated antibodies: mouse anti-human CD45-FITC (1:50, Thermo Fisher Scientific, 11-0459-42, clone [HI30]), mouse anti-mouse CD45.1-PE-Cy7 (1:100, Thermo Fisher Scientific, 25-0453-82, clone [A20]), mouse anti-human CD3-APC (1:50, Biolegend, clone [UCHT1]), mouse anti-human CD19-PE (1:50, Biolegend, 363004, clone [SJ25C1]), and mouse anti-human CD8-APC/Cy7 (1:50, Thermo Fisher Scientific, A15441, clone [RFT8]). Cells were incubated at 4°C for 30 min and washed twice with FACS Buffer and 7-AAD viability staining solution (1:50, Biolegend, 420404) was added to the cells and run with FACS Canto II (BD Biosciences). Analysis of the data was carried out using FlowJo v.10 software (Tree Star). Results are presented as mean±SEM (standard error of the mean).

### Histology and Immunofluorescence

Immediately after euthanasia by cervical dislocation, kidneys were carefully retrieved and submerged in 10% neutral buffered formalin (CellStor Pot, CellPath) for 24h hours and subsequently embedded in paraffin. 5μm serial sections were produced throughout the paraffinized kidneys. For immunostaining, slides were deparaffinized in xylene, dehydrated in graded alcohols, and rinsed with ddH2O and Tris Buffered Saline (TBS, pH 7.6). Antigen retrieval was achieved in 1mM EDTA (Sigma, E5134-500g), 10mM Tris Base (Sigma, 93352) and 0.05% Tween-20 (Sigma, P9416) buffer pH 9.0 at 96°C for 30 min, followed by 30 min cooling at room temperature. Slides were washed with TBS and blocked for 30 min with 10% Normal Donkey Serum (Abcam, ab138579) diluted in TBS containing 5% (w/v) IgG and protease-free bovine serum albumin (Jackson Immunoresearch, 001-000-162) in a humidified chamber. Primary antibodies including: anti-human CD45 (1:100, BD Biosciences, 555480), anti-human CD3 (1:100, abcam, ab11089), anti-human KRT-7 (1:200, abcam, ab68459), anti-human Ku80 (1:200, abcam, ab119935) were diluted in the blocking buffer were incubated overnight at 4°C. Secondary antibodies including: goat anti-rat IgG (H+L) Alexa Fluor 488 (Thermo Fisher Scientific, A11006), donkey anti-rabbit IgG (H+L) Alexa Fluor 555 (Thermo Fisher Scientific, A31572) donkey anti-mouse IgG (H+L) Alexa Fluor 647 (Thermo Fisher Scientific, A31571) were diluted 1:200 with DAPI-Hoechst 33342 (1:300, Invitrogen, H3570) in blocking buffer, and incubated 1 hour at room temperature. Sections were mounted in mounting medium (DAKO, S3023) under a 24×50 mm coverslip. Images were taken with an Olympus IX81 and IX73 fluorescence inverted microscope. Post-acquisition analysis was performed using ImageJ v2.0 software.

### Immunohistochemistry

Following baking at 60°C for 1hr, sections were dewaxed and rehydrated on Leica’s automated ST5020. The staining was performed on Leica’s automated Bond-III platform in conjunction with their Polymer Refine Detection System (DS9800) and a modified version of their standard template for human CD45 antibody (1:250, DAKO, M0701) using Tris EDTA (Leica’s Epitope Retrieval Solution 2, AR9640) pre-treatment at 100°C for 20 min, followed by protein block (DAKO, X090930-2), and DAB Enhancer (Leica, AR9432) for 10 mins at room temperature. Next, de-hydration and clearing were performed on Leica’s automated ST5020 and sections were mounted on Leica’s coverslipper CV5030 with mounting media DPX Mountant for Histology (Sigma Aldrich, 06522-500ML) and 22501.5 micro-coverglass coverslips (Leica Microsystems, 3800121G).

### Human CD45 Quantification

Whole slide scanning was performed on the Leica Aperio AT2 at a resolution of 0.5µm/pixel. Scanned images in the .svs file format were uploaded into HALOImage Analysis Platform version 3.6.4134 and HALO AI version 3.6.4134 (Indica Labs, Inc.). Regions for analysis were annotated manually using HALO software tools. The Multiplex IHC v3.4.9 module was used to count CD45 positive cells and CD45 positive areas. The CD45 minimum intensity for the positive threshold was set to 0.08 AU. The Area Quantification v2.4.3 module was used to measure CD45 positive pixels (area). The CD45 minimum intensity for the positive threshold was 0.1165.

### SPMC Isolation from Mouse Spleens

Spleens were harvested immediately following the killing of the animals and placed in MACS Tissue Storage Solution (Miltenyi, 130-100-008) at 4°C overnight. Tissues were dissociated into a single cell suspension by mechanical breakdown with a syringe plunger while passing through a 40µm cell strainer (FisherBrand, 22363547) in RPMI medium (Thermo Fisher Scientific, 21875-034). The resulting cell suspensions were centrifuged at 300g for 10 minutes. The pellets were resuspended in 1mL of red blood lysis buffer (StemCell Technologies, 07850), incubated for 10 mins at 4°C and wash with FACS buffer. Pellets were counted with Countess II FL (Life Technologies) and for each sample i) 5×10^5^ cells were used for flow cytometry analysis for the following conjugated antibodies: mouse anti-human CD45-FITC (1:50, Thermo Fisher Scientific, 11-0459-42, clone [HI30]), mouse anti-mouse CD45.1-PE-Cy7 (1:100, Thermo Fisher Scientific, 25-0453-82, clone [A20]) in FACS Canto II (BD Biosciences); and ii) 6×10^6^ were frozen in 10% DMSO ((Sigma, D8418-250mL) in FBS for further analysis using the Cytek 25-Color Immunoprofiling Assay, cFluor Reagent Kit (Cytek, SKU R7-40002) using Aurora spectral flow cytometry (Cytek). Analysis of the data was carried out using FlowJo v10 software (Tree Star).

### DNA sequencing

For standard whole-genome sequencing (WGS), DNA libraries were prepared using standard Illumina whole-genome library preparation protocols. For whole-genome nanorate sequencing (NanoSeq), DNA libraries were prepared as previously described in Abascal et al.^21^ For targeted NanoSeq, DNA libraries were prepared as described in Lawson et al^22^. Samples were multiplexed and sequenced on an Illumina NovaSeq 6000 platform to generate 150-base-pair (bp) paired-end reads.

### Sequence read alignment

For each sample, DNA sequence reads were aligned to the reference human genome assembly (version GRCh37/hg19) using the BWA-MEM algorithm^60^ as implemented in BWA (v0.7.17-r1188), with options ‘-T 30 -Y -p -t 8’. The aligned reads were sorted and indexed using samtools (v1.9)^61^, and duplicate reads were marked using the bammarkduplicates tool from biobambam2 (v2.0.87; gitlab.com/german.tischler/biobambam2). For NanoSeq samples, sequence reads were additionally processed using the computational workflow described by Abascal et al. (github.com/cancerit/NanoSeq).

### Variant calling and processing

For whole-genome NanoSeq samples, single-nucleotide variants (SNVs) and indels were identified using the mutation calling workflow described in Abascal et al.^21^ (v3.5.4; github.com/cancerit/NanoSeq) with default parameters. The earliest samples available from each organoid line (Line 1_PCO_P0, Line 2_PCO_Primary, Line 3_PCO_Primary) were selected as matched normal for all other samples from the same line. For targeted NanoSeq samples, single-nucleotide variants (SNVs) and indels were identified as described in Lawson et al.^22^, with the exception of an additional preprocessing step to remove reads reflecting mouse DNA contamination in these samples. Decontamination was performed as follows: a combined reference genome was produced by concatenating the human (hs37d5) and mouse (GRCm38) reference genomes, while adding a ‘mouse’ label to the name of each GRCm38 sequence; the combined reference was indexed using BWA (v0.7.17-r1188)^60^; BAM files for each sample were converted to FASTQ using samtools (v1.9)^61^ and realigned to the combined reference genome using the BWA-MEM algorithm as implemented in BWA, with options ‘-t 8 - C’; reads aligning to the mouse portion of the reference genome were removed from each realigned BAM using ‘samtools view’ followed by grep with option ‘-v $’\t’’mouse’’.

For WGS samples, SNVs were identified using the Cancer Variants through Expectation Maximization (CaVEMan) algorithm (v1.13.15)^62^, with copy number options set to ‘major=5’ and ‘minor=2’. Short insertions/deletions (indels) were identified using the Pindel algorithm (v3.3.0)^63^. CaVEMan and Pindel were run separately for each passage 1 (ePCO_P1/PCO_P1) sample, using the corresponding passage 0 sample from the wild-type organoid (PCO_P0) as the matched normal control. CaVEMan and Pindel annotate variant calls using a series of quality flags, with the ‘PASS’ flag denoting no quality issues affecting the call^62,63^. To remove false positives, calls presenting any flag other than ‘PASS’ were discarded. In addition, for SNVs, the series of post-processing filters described by Ellis et al.^64^ were applied sequentially. After filtering, SNVs were re-genotyped to obtain accurate counts of sequence reads supporting each variant allele in each sequenced sample, using the ‘bam2R’ function in the deepSNV (v1.32.0) R package^65,66^. Indels were re-genotyped using the vafCorrect tool (github.com/cancerit/vafCorrect).

### CRISPR off-target site analysis

A set of predicted CRISPR off-target sites, based on the CRISPR sgRNA sequences, was produced for the human genome (GRCh37/hg19) using the Cas-OFFinder software (v2.4)^67^ with options: PAM motif = NGG, maximum mismatches = 9, maximum DNA/RNA bulge size = 2. Predicted off-target sites were overlapped against gene CDS coordinates from Ensembl (v102; grch37.ensembl.org) and information in the COSMIC Cancer Gene Census (v98; cancer.sanger.ac.uk/census) using the GenomicRanges package (v1.42.0)^68^ in R (v4.0.2) to determine the fraction of predicted sites within cancer genes or other protein-coding genes.

Two approaches were followed to identify potential mutations near predicted off-target sites: (1) searching for high-frequency variants identified by CaVEMan or Pindel within 50 bp of any predicted site in a ePCO sample, and manually assessing the location, read support and functional consequence of identified mutations; and (2) identifying low-frequency variants *de novo* using the tools ‘mpileup’ (with option ‘-R’ used to supply a BED file containing 100-bp regions around each predicted off-target site) and ‘call’ (with options ‘-Oz -mv’) in the bcftools software (v1.9)^69^, searching for runs of ≥5 variants located within a distance of ≤20 bp, and manually inspecting these runs of variants for evidence of CRISPR-mediated editing. Visual inspection of candidate mutations was performed using the Integrative Genomics Viewer (IGV) software (v2.3.53)^70^. These analyses yielded no evidence of CRISPR-associated mutations in any ePCO sample.

### Mutation burden and rate analysis

For samples sequenced through NanoSeq, mutation burdens per diploid genome were estimated using the method described by Abascal et al.^21^ (v3.5.4; github.com/cancerit/NanoSeq). Briefly, the number of somatic mutations detected in the sample was scaled by the ratio between the number of bases sequenced to enough depth to allow base calling, and the number of bases in the diploid human genome; while also accounting for the difference in trinucleotide sequence composition between the set of sequenced bases and the reference human genome. For each organoid line, the *in vitro* mutation rate was calculated by dividing the difference in mutation burden between the earliest (P0) and latest (P1/P5) samples sequenced for that line, and the number of days elapsed between sample collection dates. For targeted NanoSeq samples, mutant cell fractions were calculated for each mutation as twice the duplex variant allele fraction (VAF) of the mutation, and then converted to mutant cell percentages for visualisation.

### Mutational signature analysis

Mutational signatures of single-nucleotide variants (single-base substitutions) on a trinucleotide sequence context were inferred from whole-genome-normalised NanoSeq mutation counts using the sigfit (v2.0) R package^32^. To obtain accurate estimates of signature exposure, signature deconvolution was performed using an approach that combines signature fitting and extraction in a single inference process^32^. More specifically, the ‘fit_extract_signatures’ function in sigfit was used to fit COSMIC signatures SBS1 and SBS5 (retrieved from the COSMIC v3.2 signature catalogue; cancer.sanger.ac.uk/signatures) to the normalised mutation counts per sample, while simultaneously extracting two additional signatures *de novo* (SBS-A and SBS-B). The inferred signatures were subsequently re-fitted to the mutational spectra of mutations in each sample (using the ‘fit_signatures’ function in sigfit) in order to estimate the exposure of each sample to each signature. This fitting process was performed separately for each sample. Signatures SBS-A and SBS-B were only included in a sample’s fitting if the lower limits of the 95% highest posterior density intervals of their respective exposures in that sample (as inferred by ‘fit_extract_signatures’) were greater than 0.01.

### Copy number analysis

For each WGS sample, allele-specific copy number (CN) was assessed using the ASCAT NGS software (v4.4.1)^71^ with default options. This method infers segments of allele-specific copy number along a genome, exploiting two sources of information from: the log-transformed sequencing coverage ratio between the target and matched normal samples (logR), and the B-allele fraction (BAF; the fraction of reads covering a germline SNP that support one of the alleles).

### Plasma Membrane Labelling

Cell surface proteins were labelled essentially as described in Weekes et al^72^. Briefly, cells are incubated with 10mM Sodium Periodate, 100µM Aminooxy Biotin and 10µM Aniline in PBS pH 6.7 for 30 minutes at 4 degrees. Cells were then lysed in 50mM TEAB pH8.5, 150mM NaCl, 1% Triton X-100 and protease inhibitor cocktail (Roche, Complete) by rotating end-over-end at 4 degrees for 30 minutes. Lysates are then cleared of nuclei by centrifugation at 13’000g for 10minutes before being added to 30µL high capacity Neutravidin-agarose beads (Thermo Fisher) and incubated for 2h with end-over-end rotation at 4 degrees. Using a postive pressure manifold (Tecan M10) beads were washed extensively with 400uL of the following: 10x lysis buffer, 10x 50mM TEAB/0.5% SDS, 10x 50mM TEAB/6M urea, 3x 50mM TEAB. After 5 urea washes, the beads were resuspended in urea wash buffer containing 10mM TCEP and 40mM Chloroacetamide and incubated with shaking for 45minutes at room temperature. After washing, beads were removed to microfuge tubes and resuspended in 50µL 50mM TEAB with 0.5ug Trypsin/LysC mix (Promega) and digested for 6h at 37 degrees with shaking at 1250rpm using a Thermomixer (Eppendorf). Supernatants were then removed from the beads and dried prior to labelling with TMT reagents.

### TMT Labelling and clean-up

Samples were resuspended in 21µL 100mM TEAB pH 8.5. After allowing to come to room temperature, 0.5mg TMT reagents (Thermo Fisher) were resuspended in 9µL anhydrous ACN which was added to the respective samples and incubated at room temperature for 1h. Sample were labelled as follows with TMT labels of increasing mass designated TMT 1-18: (3x Line 2 PCO with TMT1,3 and 5), 3x (Line 2 ePCO P1 with TMT 9, 11 and 13), (3x Line 2 ePCO P5 with TMT 10, 12 and 14). A 3µL aliquot of each sample was taken and pooled to check TMT labelling efficiency and equality of loading by LC-MS. Samples were stored at −80°C in the interim. After checking each sample was at least 96% TMT labelled total reporter ion intensities were used to normalise the pooling of the remaining samples such that the final pool should be as close to a 1:1 ratio of total peptide content between samples as possible. This final pool was brought up to a final volume of 1mL with 0.1% TFA. FA was added until the pH was <2. The sample was then cleaned up by SPE using a 50mg tC18 SepPak cartridge (Waters) and a positive pressure manifold. The cartridge was wetted with 1mL 100% Methanol followed by 1mL ACN, equilibrated with 1mL 0.1% TFA and the sample loaded slowly. The cartridge was washed 3x with 1mL 0.1% TFA before eluting sequentially with 250µL 40% ACN and 750µL 80% ACN before drying in a vacuum centrifuge.

### Basic pH reversed phase fractionation

TMT labelled samples were resuspended in 40µL 200mM Ammonium formate pH10 and transferred to a glass HPLC vial. BpH-RP fractionation was conducted on an Ultimate 3000 UHPLC system (Thermo Scientific) equipped with a 2.1 mm × 15 cm, 1.7µ Kinetex EVO column (Phenomenex). Solvent A was 3% ACN, Solvent B was 100% ACN, solvent C was 200 mM ammonium formate (pH 10). Throughout the analysis solvent C was kept at a constant 10%. The flow rate was 500 µL/min and UV was monitored at 280 nm. Samples were loaded in 90% A for 10 min before a gradient elution of 0–10% B over 10 min (curve 3), 10-34% B over 21 min (curve 5), 34-50% B over 5 mins (curve 5) followed by a 10 min wash with 90% B. 15s (100µL) fractions were collected throughout the run. Fractions containing peptide (as determined by A280) were recombined across the gradient to preserve orthogonality with online low pH RP separation. For example, fractions 1, 25, 49, 73, 97 are combined and dried in a vacuum centrifuge and stored at −20°C until LC-MS analysis. 12 Fractions were generated in this manner.

### LC-MS

Samples were analysed on an Orbitrap Fusion instrument on-line with an Ultimate 3000 RSLC nano UHPLC system (Thermo Fisher). Samples were resuspended in 10µL 5% DMSO/0.1% TFA and all sample was injected. Trapping solvent was 0.1% TFA, analytical solvent A was 0.1% FA, solvent B was ACN with 0.1% FA. Samples were loaded onto a trapping column (300µm x 5mm PepMap cartridge trap (Thermo Fisher)) at 10µL/min for 5 minutes at 60 degrees. Samples were then separated on a 75cm x 75µm i.d. 2µm particle size EasySpray PepMap C18 column (Thermo Fisher) at 60 degrees. The gradient was 3-10% B over 10mins, 10-35% B over 155 minutes, 35-45% B over 9 minutes followed by a wash at 95% B for 5minutes and requilibration at 3% B. Eluted peptides were introduced by electrospray to the MS by applying 1.5kV to the emitter. During the gradient elution, mass spectra were acquired with the following parameters: MS1 scans were acquired at 120’000 resolution and the TopN most abundant precursors in a 3s cycle time taken for MS2 and MS3. Selected precursors were fragmented by CID in the ion trap at a NCE of 30 and scanned out in the ion trap in “rapid” mode. Each MS2 scan triggered an MS3/SPS scan where the top 10 most abundant precursor ions were selected a fragmented by HCD at NCE 55. TMT reporter ions were scanned out in the Orbitrap at a resolution of 50’000. All settings for AGC target were set to automatic.

### Proteomics Data Processing

Raw data were processed with Peaks11 (Bioinfor). Tolerances for mass deviation were 15ppm MS1, 0.5Da MS2 and 0.02Da MS3. Fixed modifications were Carbamidomethylated Cysteine, and TMTpro at Peptide N-termini and Lysine. Variable modifications were Oxidised Methionine, Acetylted Protein N-termini and Deamidated Asparagine and Glutamine. The database was UniProt Homo Sapiens, reviewed entries and canonical isoforms only. An in house generated contaminant database was also used. PSM FDR was controlled at 0.1% resulting in a protein group FDR of ∼1%. Automatic normalisation based on TIC was used. Identified proteins and their TMT abundances were output to .csv, imported to R and submitted to statistical analysis using LIMMA, a moderated t-test available through the Bioconductor package^73^. LIMMA p-values were corrected for multiple hypothesis testing using the Benjamini-Hochberg method to generate an FDR (q-value) for each comparison.

### Spatial Transcriptomics

GeoMx experiments were performed in 5μm paraffin-embedded slides. The sections used in the experiments included 3 paired kidney samples of PCO and ePCOs from PBMC donor A.

#### Tissue Preparation

Tissue preparation for GeoMx experiments were performed as per the Nanostring Document Library manual MAN-10150-06 using the manual RNA Slide Preparation Protocol (FFPE). Briefly, the tissue sections were deparaffinized, rehydrated, antigen retrieved (20 minutes), RNA targets were exposed using proteinase K (15 minutes) and post fixed to preserve the tissue morphology. The slides were then in situ hybridized with GeoMx Human Whole Transcriptome Atlas NGS probes for 18 hours overnight. Next day, the probes were washed in stringent wash buffer to remove the off-target probes and blocked with buffer W (Nanostring Technologies). Morphology marker staining was done using a custom CD45 (1:100, Cell Signalling Technologies, 13917, clone [D9M8I]), PanCK (1:40, 532 excitation, Novus, NBP2-33200, clone [AE1+AE3]) and SYTO13 (1:10, 488 excitation, Nanostring Technologies). AF647 (1:200, Thermo Fisher Scientific, A31573) was used as secondary antibody for the CD45 antibody, and the slides were washed before after with 2x SCC two times 5 minutes each. Stained slides were loaded onto the GeoMx instrument and scanned for region of interest (ROI) selection.

#### ROI Selection and Data Acquisition

For GeoMx DSP sample collections, entire slides were imaged at 20x magnification and morphologic markers were used to select ROIs using organic shapes: PanCK staining was utilized to visualise cholangiocytes and CD45 staining was used to visualise immune cells. ROIs were selected by outlining custom polygonal shapes around the cholangiocyte clusters. ROIs were further segmented based on PanCK^+^ SYTO13^+^ or CD45^+^SYTO13^+^ staining conditions to select specifically the cholangiocytes or CD45+ cells rich areas within these ROIs. Advanced Parameters such as Erode, N-Dilate, Hole size, and Particle size were adjusted as seemed fit for each slide. RNA reporters from each segment were collected separately for sequencing. These segmentations defined areas of interest (AOIs) which were then exposed to 385nm light (UV), releasing the indexing oligonucleotides which were collected with a micro capillary and deposited into a 96-well plate for subsequent processing. The indexing oligonucleotides were dried overnight and resuspended in 10uL of DEPC-treated water.

#### Library preparation and sequencing

Library preparation was performed as mentioned in Nanostring Document Library MAN-10153-05-1. Specifically, sequencing libraries were generated by PCR from the photo-released indexing oligos and AOI-specific Illumina adapter sequences. Each PCR reaction used 4μl of indexing oligonucleotides, 4μl of indexing PCR primers, 2μl of Nanostring 5X PCR Master Mix. Thermocycling conditions were 37 °C for 30min, 50 °C for 10min, 95 °C for 3min; 18 cycles of 95 °C for 15s, 65 °C for 1min, 68 °C for 30s; and 68 °C for 5min. PCR reactions were pooled and AMPure bead cleanup; and pooled libraries were paired-sequenced on an Illumina Novoseq 6000 instrument (processed by Azenta).

#### Bioinformatic Analysis

FASTQ files for each sample were converted to digital count conversion (DCC) files using the GeoMx DnD pipeline (v.1) of Nanostring according to manufacturer’s pipeline. DCC files were imported back into the GeoMx DSP instrument for QC and data analyses using GeoMx DSP analysis suite version 2.2.0.111 (Nanostring). A minimum of 10,000 reads were required for each sample. Probes were checked for outlier status by implementing a global Grubb’s outlier test with alpha set to 0.01. The counts for all remaining probes for a given target were then collapsed into a single metric by taking the geometric mean of probe counts. For each sample, an RNA-probe-pool-specific negative probe normalization factor was generated on the basis of the geometric mean of negative probes in each pool. To ensure good data quality, We then calculated the 75th percentile of the gene counts (that is, geometric mean across all non-outlier probes for a given gene) for each AOI, and normalized to the geometric mean of the 75th percentile across all AOIs to give the upper quartile (Q3) normalization factors for each AOI. The distribution of these Q3 normalization factors were then checked for outliers. We subsequently removed sample quantification that are beyond the limit of quantification at 2, namely sample DSP-1001660035014-A-E05.

Cell population quantification was performed on Q3 normalized spatial transcriptomics data using SpatialDecon (ver. 1.6.0). Transcriptomics was captured across two batches and corrected using the RUV4 function at k = 10 in standR (ver. 1.8.0). Deconvolution of immune cells at CD45+ ROIs was performed with the SafeTME and Immune Census HCA dataset using SpatialDecon (ver. 1.6.0). Differential expression analysis of DKO vs WT was executed using EdgeR (ver. 3.38.4) and limma voom (ver. 3.52.4). Gene set enrichment analysis was performed with clusterprofiler (ver. 4.15.1.001) based on the calculated foldchange for either CD45+ or PanCK+ compartments against pathways in Gene Ontology (GO) at minGSSize = 3, nPermSimple = 10000, maxGSSize = 800, pvalueCutoff = 0.05, and pAdjustMethod = "BH". To allow comprehensive understanding of the key pathways, namely T cell, B cell, and cytotoxicity in CD45+ compartments and cytoxocitiy, allograft rejection, apoptosis, and stress-associated pathways in PanCK+ compartments, GO pathways containing character patterns were extracted from the GSEA results. For PanCK+ compartments, additional GSEA analyses against GO (and merged using the simplify() function by adjusted p.value at 0.7 using Wang measurement to remove redundant terms), KEGG, and Reactome were also performed and a lenient filter of p-value at 0.5 were applied to extract GO, KEGG, and Reactome pathways/terms associated to cytotoxicity, allograft rejection, apoptosis, and stress-associated pathways. Subsequently, ggplot2 (ver. 3.5.1), circlize (ver. 0.4.16), and pathview(ver. 1.36.1) are used to visualize spatial transcriptomics results.

### Bioinformatics Software

Mutation data analyses were performed using the following software: ASCAT NGS 4.4.1, bcftools 1.9, biobambam2 2.0.87, BWA-MEM 0.7.17-r1188, Cas-OFFinder 2.4, CaVEMan 1.13.15, IGV 2.3.53, NanoSeq workflow 3.5.4, Pindel 3.3.0, samtools 1.9, vafCorrect 5.4.0. Analyses of mutation data were performed on R 4.0.2, using the packages: Biostrings 2.58.0, deepSNV 1.32.0, dndscv 0.0.1.0, GenomicRanges 1.42.0, rtracklayer 1.50.0, sigfit 2.0, stringr 1.5.1. GeoMx Spatial Transcriptomics data has been performed using R 4.2.2.

### Statistical Analysis

For statistical analyses, unpaired t-tests were performed to assess differences in IFN-ɣ secretion and percentage of 7AAD-positive cells between PCO and ePCO *in vitro* co-cultures. Paired (PCO vs ePCO) or unpaired t-tests (PCO vs Matrigel-only; ePCO vs Matrigel-only) were performed to assess hCD45+ infiltration in humanised mice groups. Differentially expressed surface proteins in Line 2 ePCO compared to its parental PCO; in Line 2 ePCO passage 5 (P5) compared to PCO; and Line 2 ePCO passage 5 (P5) compared to Line 2 ePCO passage 1 (P1) were identified by q-value of <=0.01 and fold change cut off of 2-fold.

## DATA AVAILABILITY

All data are available in the main text or the supplementary materials. DNA sequencing data have been deposited on the European Genome-phenome Archive (ega-archive.org) under study accession numbers EGAS00001006326, EGAS00001006405 and EGAS00001007266.

Human mutational signatures were retrieved from the COSMIC v3.2 database (cancer.sanger.ac.uk). Custom code for mutation analyses has been deposited on GitHub (github.com/baezortega/ePCO2025). Mass spectrometry proteomics data have been deposited to the ProteomeXchange Consortium via the PRIDE^74^ partner repository with the dataset identifier PXD052302 and 10.6019/PXD052302. Spatial transcriptomics data (FASTQ files, processed count matrices, annotation files, meta data) are available on the Gene Expression Omnibus (GEO: GSE285434). Custom code for spatial transcriptomics analyses is available on GitHub (github.com/chengwailei/HLA_DKO_SpatialTranscriptomics_Analysis). Cholangiocyte organoids described in this publication are available from University of Cambridge under a material transfer agreement with the university.

## REFERENCES

1. Blau, H. M. & Daley, G. Q. Stem Cells in the Treatment of Disease. N. Engl. J. Med. 380, 1748–1760 (2019).

2. Hoang, D. M. et al. Stem cell-based therapy for human diseases. Signal Transduct Target Ther 7, 272 (2022).

3. Yamanaka, S. Pluripotent Stem Cell-Based Cell Therapy—Promise and Challenges. Cell Stem Cell 27, 523–531 (2020).

4. Method of the Year 2017: Organoids. Nat. Methods 15, 1–1 (2018).

5. Artegiani, B. & Clevers, H. Use and application of 3D-organoid technology. Hum. Mol. Genet. 27, R99–R107 (2018).

6. Huch, M., Knoblich, J. A., Lutolf, M. P. & Martinez-Arias, A. The hope and the hype of organoid research. Development 144, 938–941 (2017).

7. Kim, J., Koo, B.-K. & Knoblich, J. A. Human organoids: model systems for human biology and medicine. Nat. Rev. Mol. Cell Biol. 21, 571–584 (2020).

8. Shankaran, A., Prasad, K., Chaudhari, S., Brand, A. & Satyamoorthy, K. Advances in development and application of human organoids. 3 Biotech 11, 257 (2021).

9. Tang, X.-Y. et al. Human organoids in basic research and clinical applications. Signal Transduct Target Ther 7, 168 (2022).

10. Petrus-Reurer, S. et al. Immunological considerations and challenges for regenerative cellular therapies. Commun Biol 4, 798 (2021).

11. Kim, J. J., Fuggle, S. V. & Marks, S. D. Does HLA matching matter in the modern era of renal transplantation? Pediatr. Nephrol. 36, 31–40 (2021).

12. Zachary, A. A. & Leffell, M. S. HLA Mismatching Strategies for Solid Organ Transplantation - A Balancing Act. Front. Immunol. 7, 575 (2016).

13. Hafeez, M. S. et al. HLA mismatch is important for 20-year graft survival in kidney transplant patients. Transpl. Immunol. 80, 101861 (2023).

14. Koga, K., Wang, B. & Kaneko, S. Current status and future perspectives of HLA-edited induced pluripotent stem cells. Inflamm. Regen. 40, 23 (2020).

15. Höijer, I. et al. CRISPR-Cas9 induces large structural variants at on-target and off-target sites in vivo that segregate across generations. Nat. Commun. 13, 627 (2022).

16. Guo, C., Ma, X., Gao, F. & Guo, Y. Off-target effects in CRISPR/Cas9 gene editing. Front. Bioeng. Biotechnol. 11, 1143157 (2023).

17. Lopes, R. & Prasad, M. K. Beyond the promise: evaluating and mitigating off-target effects in CRISPR gene editing for safer therapeutics. Front. Bioeng. Biotechnol. 11, 1339189 (2023).

18. Kuijk, E. et al. The mutational impact of culturing human pluripotent and adult stem cells. Nat. Commun. 11, 1–12 (2020).

19. Merkle, F. T. et al. Human pluripotent stem cells recurrently acquire and expand dominant negative P53 mutations. Nature 545, 229–233 (2017).

20. Rouhani, F. J. et al. Substantial somatic genomic variation and selection for BCOR mutations in human induced pluripotent stem cells. Nat. Genet. 54, 1406–1416 (2022).

21. Abascal, F. et al. Somatic mutation landscapes at single-molecule resolution. Nature 593, 405–410 (2021).

22. Lawson, A. R. J. et al. Somatic mutation and selection at epidemiological scale. medRxiv 2024.10.30.24316422 (2024) doi:10.1101/2024.10.30.24316422.

23. Sampaziotis, F. et al. Reconstruction of the mouse extrahepatic biliary tree using primary human extrahepatic cholangiocyte organoids. Nat. Med. 23, 954–963 (2017).

24. Tysoe, O. C. et al. Isolation and propagation of primary human cholangiocyte organoids for the generation of bioengineered biliary tissue. Nat. Protoc. 14, 1884–1925 (2019).

25. Sampaziotis, F. et al. Cholangiocyte organoids can repair bile ducts after transplantation in the human liver. Science 371, 839–846 (2021).

26. Sondka, Z. et al. The COSMIC Cancer Gene Census: describing genetic dysfunction across all human cancers. Nat. Rev. Cancer 18, 696–705 (2018).

27. Ben-David, U. et al. Aneuploidy induces profound changes in gene expression, proliferation and tumorigenicity of human pluripotent stem cells. Nat. Commun. 5, 4825 (2014).

28. Ben-David, U. & Benvenisty, N. High prevalence of evolutionarily conserved and species-specific genomic aberrations in mouse pluripotent stem cells. Stem Cells 30, 612–622 (2012).

29. Broberg, K., Toksvig-Larsen, S., Lindstrand, A. & Mertens, F. Trisomy 7 accumulates with age in solid tumors and non-neoplastic synovia. Genes Chromosomes Cancer 30, 310– 315 (2001).

30. Zhang, X.-H., Tee, L. Y., Wang, X.-G., Huang, Q.-S. & Yang, S.-H. Off-target effects in CRISPR/Cas9-mediated genome engineering. Mol. Ther. Nucleic Acids 4, e264 (2015).

31. Rasul, M. F. et al. Strategies to overcome the main challenges of the use of CRISPR/Cas9 as a replacement for cancer therapy. Mol. Cancer 21, 64 (2022).

32. Gori, K. & Baez-Ortega, A. sigfit: flexible Bayesian inference of mutational signatures. bioRxiv 372896 (2020) doi:10.1101/372896.

33. Alexandrov, L. B. & Stratton, M. R. Mutational signatures: the patterns of somatic mutations hidden in cancer genomes. Curr. Opin. Genet. Dev. 24, 52–60 (2014).

34. Alexandrov, L. B. et al. The repertoire of mutational signatures in human cancer. Nature 578, 94–101 (2020).

35. Behjati, S. et al. Genome sequencing of normal cells reveals developmental lineages and mutational processes. Nature 513, 422–425 (2014).

36. Rouhani, F. J. et al. Mutational History of a Human Cell Lineage from Somatic to Induced Pluripotent Stem Cells. PLoS Genet. 12, e1005932 (2016).

37. Zou, X. et al. Validating the concept of mutational signatures with isogenic cell models. Nat. Commun. 9, 1744 (2018).

38. Kärre, K., Ljunggren, H. G., Piontek, G. & Kiessling, R. Selective rejection of H-2-deficient lymphoma variants suggests alternative immune defence strategy. Nature 319, 675–678 (1986).

39. Ljunggren, H. G. & Karre, K. In search of the “missing self”: MHC molecules and NK cell recognition. Immunol. Today 11, 237–244 (1990).

40. Geurts, M. H. & Clevers, H. CRISPR engineering in organoids for gene repair and disease modelling. Nature Reviews Bioengineering 1, 32–45 (2023).

41. Skoufou-Papoutsaki, N. et al. Efficient genetic editing of human intestinal organoids using ribonucleoprotein-based CRISPR. Dis. Model. Mech. 16, (2023).

42. Artegiani, B. et al. Probing the Tumor Suppressor Function of BAP1 in CRISPR-Engineered Human Liver Organoids. Cell Stem Cell 24, 927–943.e6 (2019).

43. Okamoto, T., Natsume, Y., Yamanaka, H., Fukuda, M. & Yao, R. A protocol for efficient CRISPR-Cas9-mediated knock-in in colorectal cancer patient-derived organoids. STAR Protoc 2, 100780 (2021).

44. Takeda, H. et al. CRISPR-Cas9–mediated gene knockout in intestinal tumor organoids provides functional validation for colorectal cancer driver genes. Proceedings of the National Academy of Sciences 116, 15635–15644 (2019).

45. Fujii, M., Clevers, H. & Sato, T. Modeling Human Digestive Diseases With CRISPR-Cas9– Modified Organoids. Gastroenterology 156, 562–576 (2019).

46. Sun, D. et al. A functional genetic toolbox for human tissue-derived organoids. Elife 10, (2021).

47. Schwank, G. et al. Functional repair of CFTR by CRISPR/Cas9 in intestinal stem cell organoids of cystic fibrosis patients. Cell Stem Cell 13, 653–658 (2013).

48. Driehuis, E. & Clevers, H. CRISPR/Cas 9 genome editing and its applications in organoids. Am. J. Physiol. Gastrointest. Liver Physiol. 312, G257–G265 (2017).

49. Hendriks, D., Artegiani, B., Hu, H., Chuva de Sousa Lopes, S. & Clevers, H. Establishment of human fetal hepatocyte organoids and CRISPR-Cas9-based gene knockin and knockout in organoid cultures from human liver. Nat. Protoc. 16, 182–217 (2021).

50. Yoshihara, E. et al. Immune-evasive human islet-like organoids ameliorate diabetes. Nature (2020) doi:10.1038/s41586-020-2631-z.

51. Hu, X. et al. Human hypoimmune primary pancreatic islets avoid rejection and autoimmunity and alleviate diabetes in allogeneic humanized mice. Sci. Transl. Med. 15, eadg5794 (2023).

52. Deuse, T. et al. Hypoimmunogenic derivatives of induced pluripotent stem cells evade immune rejection in fully immunocompetent allogeneic recipients. Nat. Biotechnol. 37, 252–258 (2019).

53. Gornalusse, G. G. et al. HLA-E-expressing pluripotent stem cells escape allogeneic responses and lysis by NK cells. Nat. Biotechnol. 35, 765–772 (2017).

54. Han, X. et al. Generation of hypoimmunogenic human pluripotent stem cells. Proceedings of the National Academy of Sciences 116, 10441–10446 (2019).

55. Hu, X. et al. Hypoimmune induced pluripotent stem cells survive long term in fully immunocompetent, allogeneic rhesus macaques. Nat. Biotechnol. 42, 413–423 (2023).

56. Abou-Khalil, R. & Colnot, C. Renal capsule transplantations to assay skeletal angiogenesis. Methods Mol. Biol. 1130, 99–110 (2014).

57. Park, H.-C. et al. Renal capsule as a stem cell niche. Am. J. Physiol. Renal Physiol. 298, F1254–62 (2010).

58. Meissner, T. B., Schulze, H. S. & Dale, S. M. Immune Editing: Overcoming Immune Barriers in Stem Cell Transplantation. Curr Stem Cell Rep 8, 206–218 (2022).

59. Ye, Q. et al. Generation of universal and hypoimmunogenic human pluripotent stem cells. Cell Prolif. 53, e12946 (2020).

60. Li, H. Aligning sequence reads, clone sequences and assembly con*gs with BWA-MEM. (2014) doi:10.6084/M9.FIGSHARE.963153.V1.

61. Li, H. et al. The Sequence Alignment/Map format and SAMtools. Bioinformatics 25, 2078– 2079 (2009).

62. Jones, D. et al. cgpCaVEManWrapper: Simple Execution of CaVEMan in Order to Detect Somatic Single Nucleotide Variants in NGS Data. Curr. Protoc. Bioinformatics 56, 15.10.1–15.10.18 (2016).

63. Raine, K. M. et al. cgpPindel: Identifying Somatically Acquired Insertion and Deletion Events from Paired End Sequencing. Curr. Protoc. Bioinformatics 52, 15.7.1–15.7.12 (2015).

64. Ellis, P. et al. Reliable detection of somatic mutations in solid tissues by laser-capture microdissection and low-input DNA sequencing. Nat. Protoc. 16, 841–871 (2021).

65. Gerstung, M. et al. Reliable detection of subclonal single-nucleotide variants in tumour cell populations. Nat. Commun. 3, 811 (2012).

66. Gerstung, M., Papaemmanuil, E. & Campbell, P. J. Subclonal variant calling with multiple samples and prior knowledge. Bioinformatics 30, 1198–1204 (2014).

67. Bae, S., Park, J. & Kim, J. S. Cas-OFFinder: a fast and versatile algorithm that searches for potential off-target sites of Cas9 RNA-guided endonucleases. Bioinformatics 30, 1473– 1475 (2014).

68. Lawrence, M. et al. Software for computing and annotating genomic ranges. PLoS Comput. Biol. 9, e1003118 (2013).

69. Danecek, P. et al. Twelve years of SAMtools and BCFtools. Gigascience 10, (2021).

70. Thorvaldsdóttir, H., Robinson, J. T. & Mesirov, J. P. Integrative Genomics Viewer (IGV): high-performance genomics data visualization and exploration. Brief. Bioinform. 14, 178– 192 (2013).

71. Raine, K. M. et al. ascatNgs: Identifying Somatically Acquired Copy-Number Alterations from Whole-Genome Sequencing Data. Curr. Protoc. Bioinformatics 56, 15.9.1–15.9.17 (2016).

72. Weekes, M. P. et al. Proteomic plasma membrane profiling reveals an essential role for gp96 in the cell surface expression of LDLR family members, including the LDL receptor and LRP6. J. Proteome Res. 11, 1475–1484 (2012).

73. Ritchie, M. E. et al. limma powers differential expression analyses for RNA-sequencing and microarray studies. Nucleic Acids Res. 43, e47 (2015).

74. Perez-Riverol, Y. et al. The PRIDE database resources in 2022: a hub for mass spectrometry-based proteomics evidences. Nucleic Acids Res. 50, D543–D552 (2022).

